# Recalled parental bonding interacts with oxytocin receptor gene polymorphism in modulating anxiety and avoidance in adult relationships

**DOI:** 10.1101/2020.07.01.181644

**Authors:** Ilaria Cataldo, Andrea Bonassi, Bruno Lepri, Jia Nee Foo, Peipei Setoh, Gianluca Esposito

**Affiliations:** Department of Psychology and Cognitive Science, University of Trento, Rovereto, Italy; Mobile and Social Computing Lab, Fondazione Bruno Kessler, Trento, Italy; Lee Kong Chian School of Medicine, Nanyang Technological University, Singapore, Singapore; Human Genetics, Genome Institute of Singapore, Singapore, Singapore; Psychology Program, School of Social Sciences, Nanyang Technological University, Singapore, Singapore

**Keywords:** Parental Bonding, oxytocin receptor gene, gene-environment interaction, rs53576

## Abstract

Early interactions with significant individuals affect social experience throughout the course of a lifetime, as a repeated and prolonged perception of different levels of care, independence or control influences the modulation of emotional regulatory processes. As many factors play a role in shaping the expectations and features of social interaction, in this study we consider the influence of parental bonding and genetic allelic variation of oxytocin receptor gene polymorphism (rs53576) over levels of experienced anxiety and avoidance in 313 young adults belonging to two different cultural contexts, namely Italy and Singapore. Results highlight a major effect of maternal characteristics, care and overprotection, with differences between the two cultural groups. Additionally, the interaction between rs53576 and maternal overprotection suggest different environmental susceptibility in the Italian sample and the Singaporean one. Implication in clinical work and future steps are described in the conclusion.

## 1. Introduction

Fundamental works in disciplines related to developmental psychology have demonstrated that early social interactions with caregivers influence different aspects of child development, such as social relationships [8, 52, 78], academic performance [66], response to stress [25, 31], individual well-being, and risk for psychopathology [54, 60, 65, 58]. In addition, emotion-related skills in children like interpretation, regulation, and communication are built on the emotional and social relationships they share with their close ones [69]. Therefore, it is likely that positive parenting practices, like higher levels of warmth, care, and sensitivity, will result in more adaptive emotion-regulation skills [46]. The quality of parental bonding influences the development of top-down processes of emotional regulation, which could potentially shape social interactions and physiological responses in individuals. The ability to perceive typical social situations as nonthreatening and risk-free also has implications for the autonomic nervous system [55]. According to the literature, ideal parental practices can be described in terms of higher levels of care and lower overprotection. In fact, care allows individuals to be less restrained in social interactions, while overprotection diminishes the effectiveness of emotional regulation and the attitude about exploring new social situations. As a consequence, less positive caregiving patterns are also relevant to the manifestation of anxiety in social relationships [61].

According to the original theories on attachment by Bowlby and Ainsworth, attachment-specific features were considered to extend beyond cultural boundaries or parenting practices; in other words, Bowlby’s and Ainsworth’s original theory was considered to be culturally universal in the sense that parental behavior that is sensitive to the child’s needs will result in attachment security across a diverse array of cultural and ecological contexts.

Despite this assumption, the ecological components of attachment have always been seen as an important factor that needs to be taken into account, together with other features that might play a role in child-rearing behavior, such as socioeconomic status or ethnicity. In fact, one of the major concerns in the attachment-related research fields is the limitation due to results coming primarily from countries belonging to Western societies. With regards to cultural studies, Ainsworth’s canonical work in Uganda and Baltimore underlined that processes related to maternal behavior reflect specific universal features that cross cultural, geographic, and linguistic boundaries [1]. In their research based on a sample of Japanese mothers, Behrens and colleagues found that maternal attachment was predictive of the child’s attachment classification with a distribution comparable to global results [4]. In a review on parental sensitivity in ethnic minority families, Mesman highlighted the positive role of a secure pattern of attachment on children’s development [45]. Central to the quality of parenting practices and attachment are the dimensions of care and protection—encompassed in the construct of parental bonding [52]. Parental care refers to the sensitivity of parents towards the child’s needs (emotional, physical, psychological), while parental overprotection refers to the excessive restrictions (emotional, physical, psychological) imposed on the child. Partially inconsistent results have been described in a recent work on the link between the caregiver’s warmth and levels of anxiety and avoidance in adult attachment in a Chinese population [71].

### 1.1. Familial Bonds in Italy and Singapore

In light of these considerations, it appears clear that, to have a proper understanding of parental bonds, it is essential to take a wider look at the concept of the familial nucleus according to the country where these family units are immersed. In the current study, we focused on two countries, one more Western oriented (Italy) and another one representative of a non-Western model (Singapore). In Italy, which is configured within individualistic countries, family is seen as an expanded network of relationships and ties, where much attention is given to the child. The relationship that forms between parent and child tends to be supportive and encouraging towards independence, presenting high levels of emotional bonding [40]. On the other hand, Singapore is described as a multicultural context, comprised of a majority of Chinese, Indian, and Malay families. In Singapore, which belongs to a collectivist culture, family is considered the first group a person joins in his/her life. The family frame belongs to a collectivist scaffolding, like other Asian countries, where interdependent relationships occur on a social and familiar level, especially for Chinese and Indian households. However, for all ethnicities, the extended family traditionally used to live with the original family, and parental control over the children was extended until adulthood. With the sudden rise of technology and globalization, it is difficult, for the Singaporeans, to maintain a multigenerational structure in families, and many people decide either to live alone or not to have children. Furthermore, parental roles change for those who decide to have babies, especially in the last couple of decades: if the hierarchy once used to be based on a patriarchal model, with the father having disciplinary power and the mother staying at home taking care of family duties, now both parents share the care of their children. Although evidence coming from different cultures and ethnicities has been reported and discussed, concerns about the application of Western-based methodologies to non-Western environments still remain. In fact, attachment-related mechanisms are, in turn, influenced by emotional communication, and the way emotions are expressed and experienced notably differs from country to country [34]. Furthermore, while attachment dynamics have been profusely explored, little is known about the effect of processes that elapse between childhood and early adulthood, in the manifestation of dynamics in adult relationships.

### 1.2. OXTR Polymorphism rs53576

Considering the development of specific social characteristics as a result of the interaction between biological and environmental factors, it appears clear that behavioral facets might present a combination of genetic, environmental, and neurobiological factors. This involves the genetic regulation of neurotransmitters that are notably involved in the modulation of social functioning, such as the oxytocin receptor gene (*OXTR*) [14, 75, 73]. Evidence in the literature reports results about the involvement of oxytocin receptor gene single-nucleotide polymorphisms (SNPs) in modulating the response to social stressors [23, 80, 17]. Within the polymorphic region of *OXTR*, allelic variations of the single-nucleotide polymorphism rs53576 have been shown to be associated with different social behaviors. Specifically, the G allele appears to be linked with more optimal social development, which is described through higher levels of dispositional empathy [59, 67], favorable prosocial features [74, 64, 59, 73], and greater autonomic reactivity to social stressors [48]. On the other hand, A/A homozygotes have shown overall poorer social traits [72, 76], reduced empathy accuracy [59], and positive affect [42], all representing a less flexible social development. Although many results highlight the association between specific variations of the genotype and their determined social features, there are still some discrepancies in the literature [2, 41, 3]. With specific regards to the interaction with perceived parental warmth, higher levels of paternal care have been found to be associated with an increased heart rate response towards social cues for individuals with a G/G variation in an Italian sample [74]. A study by Kim and colleagues compared a Korean and an American population to investigate the interaction between perceived stress and *OXTR rs53576* variations in modulating the seeking of emotional support, suggesting that, among Americans, G-carriers were seeking more emotional social support compared to A/A homozygotes, while the same difference was not shown in the Korean sample. Moreover, they also compared Koreans raised in America and Koreans raised and living in Korea, finding greater emotional social support-seeking behavior in the first sample, hence highlighting the relevance of the cultural environment [37].

### 1.3. Current Study: Hypotheses

The wider purpose of the current study was to deepen our understanding of how the interaction between individual genetic features and the perception of parental warmth during childhood affects levels of anxiety and avoidance in adult relationships. The aim was to analyze the differences occurring in two different countries, like Italy and Singapore. The choice of these two contexts allowed a comparison of a Western country and a non-Western country, as environmental factors, such as the country one belongs to, may affect, on the one hand, the perception of parental features in childhood and, on the other hand, the formation of the behavioral constructs that drive social relationships. In order to explore parents’ transmitted behavior towards the child (assessed in terms of warmth/care and overprotection), we used the Parental Bonding Instrument (PBI) by Parker and colleagues [52]. To measure the levels of anxiety and avoidance in adult relationships, we employed the revised version of the Experience in Close Relationships (ECR-R) [24]. For the Singaporean sample, we used the original versions in English, while for the Italian sample, participants completed the translated and validated Italian versions [62, 11]. We hence set two hypotheses on the Western (i.e., Italian) and Eastern (i.e., Singaporean) sample, respectively, and independently of the cultural differences.

#### HP1 *Gene*\**environment interaction on the Western sample*

We expected to find a statistically significant effect of the interaction between *OXTR* polymorphism and parental bonding features (assessed using the PBI scales) over the main features of adult social relationships (measured with the ECR-R), with Italian A-carriers showing different rates of distress in terms of anxiety and avoidance than G/G homozygotes when they reported low levels of parental care and high levels of parental overprotection;

#### HP2 *Gene*\**environment interaction on the Eastern sample*

We expected to observe a statistically significant effect of the interaction between *OXTR* polymorphism and parental bonding features (assessed using the PBI scales) over the main features of adult social relationships (measured with the ECR-R), with Singaporean G-carriers exhibiting different rates of distress in terms of anxiety and avoidance than A/A homozygotes when they recalled low levels of parental care and high levels of parental overprotection.

The predicted gene*environment interactions on the two separated samples could not however disclose a distinct pattern of adult attachment between Eastern and Western samples. Given the complexity of the multiple relationship between genetic factors, cultural influences, and early parental bonding on adult attachment, potential gene*culture interactions were finally hypothesized at an observational level:

#### HP3 *Culture*\**gene*\**environment interactions on the total sample*

We predicted a differential susceptibility of *OXTR rs53576* to perceived parental bonding (assessed with the PBI subscales) between the Western (i.e., Italian) and the Eastern (i.e., Singaporean) groups in explaining the levels of anxiety and avoidance (measured using the ECR index.

### 2. Materials and Methods

The research was approved by the Ethical Committee of Nanyang Technological University (IRB-2015-08-020-01) and of the University of Trento (Prot-2017-019). Informed consent was obtained from all participants, and the study was conducted following the Declaration of Helsinki. Participants were recruited through a social media platform (students’ Facebook groups, for the Italian sample) and a web-based software research study participation and management system (Sona System, for the Singaporean sample) among non-parent students of the University of Trento, Italy, and Nanyang Technological University, Singapore. After informed consent was given, the online form of the questionnaires was sent through email to the participants (links to Qualtrics for the Singaporean sample and links to Google Modules for the Italian sample). Participants who gave consent for the genetic part of the experiment were contacted to set an appointment to obtain a buccal mucosa sample for genetic information. The genetic assessment was conducted on anonymized biosamples at the Nanyang Technological University (Singapore) and at the Department of Neurobiology and Behavior at Nagasaki University (Japan). Results from the questionnaires were anonymized at the beginning of the collection. The final group consisted of 313 participants that fully completed the two questionnaires and provided the experimenters with a DNA sample (Italian sample N = 97 (M = 38; F = 59); mean age = 23.01 (M = 23.58; F = 22.64), SD = 3.82 [range 18 ∼ 35]; Singaporean sample N = 216; M = 73; F = 143; mean age = 21.51, SD = 1.83 [range 18 ∼ 31]).

### 2.1. Questionnaires

#### Parental Bonding Instrument

The Parental Bonding Instrument (PBI) is a 50-item self-report questionnaire that investigates participants’ perception of both maternal (25 items) and paternal (25 items) care and protection in their first 16 years of life. It was developed by Parker, Tupling, and Brown in 1979 using factor analysis from parents’ self-reporting their childhood experiences, the results of which yielded two factors: warmth/care (hereafter referred to as care) and overprotection [52]. The care scale (PBI_Care) measures the degree to which a mother or a father was empathetic and caring versus cold and indifferent. The overprotection scale (PBI_OverP) measures the extent to which a parent was intrusive or, conversely, fostered independence in the subject. The measure has been shown to have high reliability, stability over time, and no association with social desirability, neuroticism, or extroversion [50].

#### Experience in Close Relationships-Revised

In order to evaluate anxiety and avoidance levels in close relationships, the Experience in Close RelationshipsRevised was employed (ECR-R; [24, 11]). The 36-item self-report questionnaire assesses two major dimensions of an individual’s attachment style (anxiety and avoidance) in a sentimental relationship. Both the anxiety and avoidance subscales consisted of 18 items each, rated on a 7-point Likert scale ranging from 1 (strongly disagree) to 7 (strongly agree). The anxiety dimension measures insecurity, jealousy, and fear of abandonment as opposed to feeling secure about the availability and responsiveness of romantic partners. The avoidance dimension measures the feeling of discomfort being close to others and tendency to refrain from attachment. Instead of assigning participants to an attachment style category, this scale yields two separate dimension scores for each participant.

### 2.2. Genotyping

This study adopted the same DNA derivation and genotyping procedure used by Bonassi and colleagues [7], using ACGT, Inc. (Wheeling, IL). DNA was extracted from each kit using the Oragene DNA purifying reagent, then concentrations were assessed through spectroscopy (NanoDrop Technologies, USA). Concentrations for each sample were magnified through polymerase chain reaction (PCR) for the *OXTR* gene *rs53576* region target. The forward and reverse primers that were used were 5-GCC CAC CAT GCT CTC CAC ATC-3 and 5-GCT GGA CTC AGG AGG AAT AGG GAC-3. For this DNA region, the frequency for the allelic distributions (A/A, A/G, G/G) differ among populations belonging to different ethnic groups. Although a unique consensus has still not been reached and may depend on the alleles’ distributions within a specific sample, the A allele is usually combined with the same heterozygous group A/G in the Western samples, whereas the G allele is more commonly paired with the heterozygous group A/G in the Eastern samples [77, 2, 74, 15]. In line with the existing literature, the present study followed the same paired allelic in the two different populations. In cross-cultural investigations analyzing different ethnic groups as a whole, reducing the number of genetic groups, A/A is maintained in contrast with the combined couples A/G and G/G [37, 12].

Here, Western participants (e.g., all Italian native participants with Caucasian ethnicity) having at least one A allele (A/A homozygotes or A/G) were classified into a single A-carriers group. Complementary to this, Eastern participants (e.g., all Singaporean native participants with Southeast-Asian ethnicity, namely 184 Chinese, 15 Malays, 8 Indian, 3 Caucasian, 3 Filipino, 1 Vietnamese, 1 Gurkha, and 1 Korean) having at least one G allele (G/G homozygotes or A/G) were treated as a single G-carriers group.

The averaged distribution of the different genotypes in the Caucasian population was 25–35% for A/A homozygotes and 65–75% for G-carriers (1000 Genomes project, ensembl.org), whereas the distribution in the Italian sample was 15% for A/A homozygotes and 85% for G-carriers. Specifically, genotype frequencies were as follows: A/A = 14 (14.43%), A/G = 35 (36.08%), G/G = 48 (49.49%). In the present Italian sample, the less frequent homozygous group A/A was merged with the heterozygous group A/G (A-carriers = 51%;

G/G = 49%) before the analysis. Participants’ age (*t* = 1.761, *p* = 0.383) and sex *X*^2^(1) = 0.016, *p* = 0.899) did not significantly differ between the G/G and A groups (see Table S1 of the Supplementary Materials). As for the Singaporean sample, the allelic distribution in the East-Asian population was 65–75% for A/A homozygotes and 25–35% for G-carriers (1000 Genomes project, ensembl.org), whereas in our sample, the distributions were 37% for A/A homozygotes and 63% for G-carriers. Specifically, genotype frequencies were: A/A = 80 (37.04%), A/G = 94 (43.52%), and G/G = 42 (19.44%). Participants’ age (*t* = 0.481, *p* = 0.631) and sex *X*^2^(1) = 0.189, *p* = 0.663) did not significantly differ between the A/A and G groups (see Table S1 of the Supplementary Materials).

Alleles’ frequency distribution and Hardy–Weinberg equilibrium parameters are reported in Table 1. In the total sample, the distribution of the alleles was as follows: A/A = 94 (30.03%), A/G = 129 (41.21%), and G/G = 90 (28.75%). Since the genotype frequencies for the whole sample (total sample = Italian sample + Singaporean sample) did not satisfy the Hardy–Weinberg equilibrium, analyses focused on the three distinct allelic combinations within the frame of an exploratory approach. Participants’ sex (*X*^2^(2) = 0.008, *p* = 0.996) did not significantly differ between the three groups. Participants’ age did not significantly differ between A/A and A/G (*t* = −0.814, *p* = 0.883), A/A and G/G (*t* = 0.417, *p* = 0.413), and G/G and A/G (*t* = −0.991, *p* = 0.323) (see Table S1 of the Supplementary Materials).

**Table 1:** Hardy–Weinberg equilibrium results for *OXTR rs53576* in the Italian sample, in the Singaporean sample, and in the total sample. The percentage frequency of each genetic group is reported between parentheses. ** *p* < 0.01.

| Genotype | A/A | A/G | G/G | $X^2$ | $p$-Value |
| --- | --- | --- | --- | --- | --- |
| rs53576<sub>ITA</sub> | 14 (14.43%) | 35 (36.08%) | 48 (49.49%) | 3.05 | 0.08 |
| rs53576<sub>SNG</sub> | 80 (37.04%) | 94 (43.52%) | 42 (19.44%) | 2.24 | 0.13 |
| rs53576<sub>Total</sub> | 94 (30.03%) | 129 (41.21%) | 90 (28–75%) | 9.64 | 0.002 ** |

### 2.3. Statistical Analyses

Data were analyzed with R (Version 4.0.0). Statistical tests were differently planned for hypothesis-driven and exploratory analysis. The hypothesis-driven tests were performed separately on the Italian and the Singaporean sample, whereas the exploratory tests were conducted on the total number of participants (total sample = Italian sample + Singaporean sample).

Overall, the statistical analyses followed a common procedure of variables’ evaluation and tests’ application.

#### 2.3.1. Preliminary Analyses

Before proceeding with the data analysis, we tested the internal validity of each questionnaire in both groups (Italy, Singapore). Preliminary tests were made in each population group to exclude any significant effect on the ECR-R-dependent variables that could be attributed to participants’ sex and age for the exploratory analysis (see Figures S1 and S2 of the Supplementary Materials). Univariate and multivariate distributions of the PBI and ECR-R scores were inspected for normality and the presence of extreme values. Seventy-six observations (Italian sample = 27; Singaporean sample = 49) were found as having a value higher/lower than 2 *SDs*above/below the mean. Any artificial restriction (i.e., outliers’ removal or treatment) was avoided to preserve the meaning associated with the scoring of each extreme observation. A high reliability of the subscales (see Table 2) also excluded potential biases due to extreme scores [79].

**Table 2:**
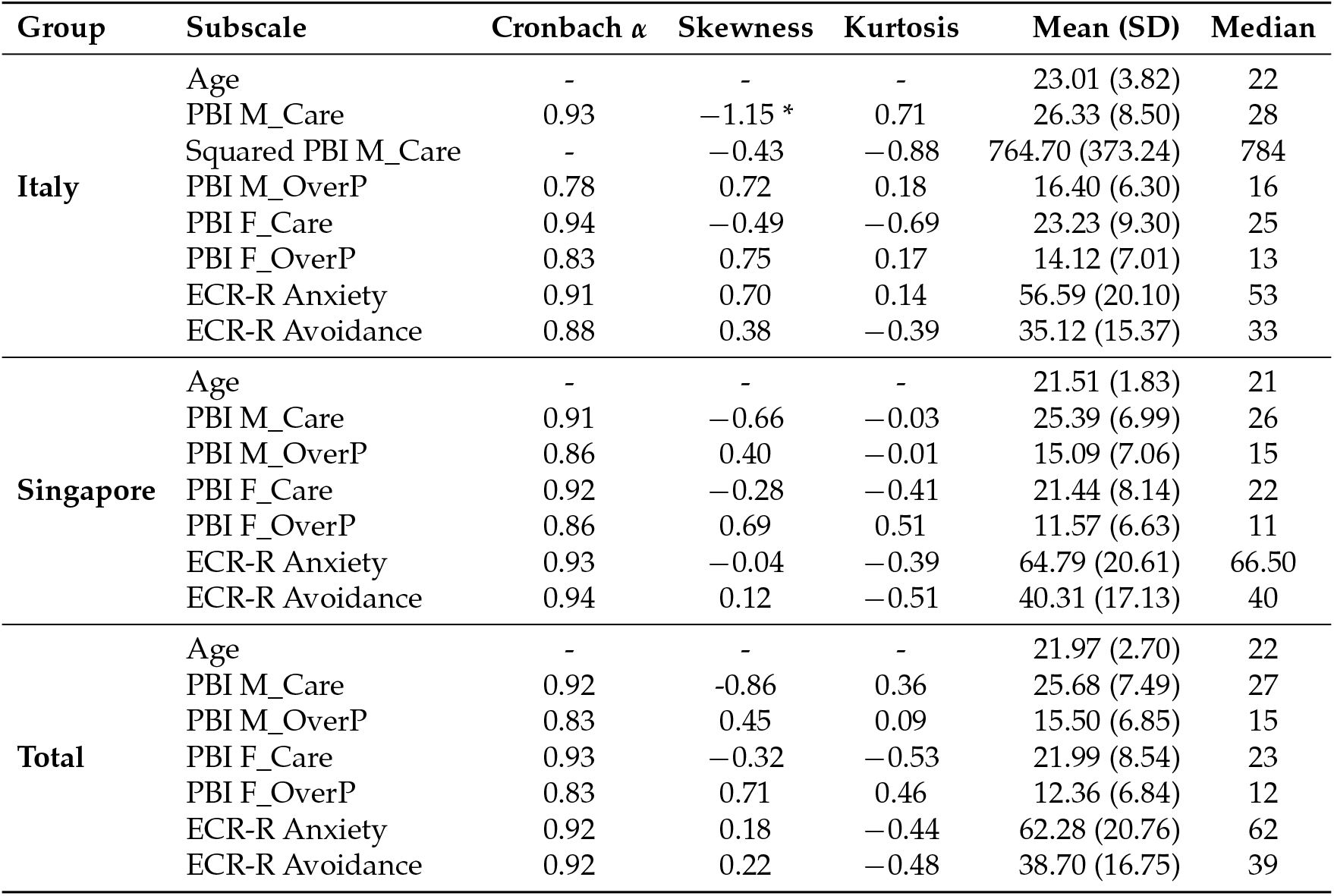
Summary of Cronbach’s *α*, skewness, and kurtosis for each subscale of the Parental Bonding Instrument and Experience in Close Relationships in the Italian sample, Singaporean sample, and total sample. Statistics of age as the continuous variable are also reported. * = squared transformation applied. M_Care and M_OverP refer to the maternal dimensions, whereas F_Care and F_OverP are related to the father.

#### 2.3.2. Main Analyses

Exploratory correlation matrices were generated for the Italian, Singaporean, and total samples to probe possible associations among the continuous variables (e.g., PBI and ECR-R dimensions).

A series of hierarchical multiple regression (HMR) analyses was performed on the Italian and Singaporean sample separately to test the hypothesis that gene*environment interaction would predict the dependent variable (DV) adult anxiety (e.g., one series on the Italian sample, one series on the Singaporean sample). A series of HMR analyses was also conducted on the Italian and Singaporean sample separately to test the hypothesis that the gene*environment interaction would predict the DV adult avoidance (e.g., one series on the Italian sample, one series on the Singaporean sample). In summary, a six-step HMR was conducted for each ECR-R subscale (namely, anxiety and avoidance) as the DV in reference to each sample (see the models applied to each step in Table S2 of the Supplementary Materials). Allelic variation for *OXTR* was entered at Stage 1 as a between-subjects factor of the regression to control for differences due to the genetics (Italian sample: G/G vs. A-carriers; Singaporean sample: A/A vs. G-carriers). Progressively, all the PBI subscales as continuous predictors were entered into each following step; specifically, maternal care was entered at Step 2, maternal overprotection at Step 3, paternal care at Step 4, and paternal overprotection at Step 5. In the final model (Step 6), the interaction between *OXTR* polymorphism variants and perceived parental bonding was tested.

A further series of HMR analyses was conducted on the total sample to test the exploratory hypothesis that culture would modulate the interaction between *OXTR* and early caregiving behavior on adult anxiety and avoidance as a unique DV (e.g., one series on the total sample for anxiety as the DV, one series on the total sample for avoidance as the DV). Therefore, in the total sample, an initial step was added in order to include the cultural variable (Italian vs. Singaporean) in the predictors of the regression model, for a total number of seven steps (see Table S2 of the Supplementary Materials). Allelic groups for *OXTR* were here added at Stage 2 as a between-subjects factors (Total sample: A/A vs. A/G vs. G/G). Successively, all the PBI dimensions were singularly entered into each subsequent step following the order of administration of each sub-dimension of the PBI questionnaire. The final model (Step 7) was here defined by the interaction between culture, *OXTR* polymorphisms variants, and perceived parental bonding.

To compare the significance of each progressive model, the analysis of variance (ANOVA) was used to analyze the contribution of each predictor added at every step. As the contrast coding system for the categorical variables (e.g., culture and *OXTR* as factors) entered into the linear models, dummy coding was applied to compare each level of a given categorical variable to a fixed reference level. The factor *OXTR* presented two levels for the analysis on the Italian (reference level: A-carriers as 1; G/G as 2) and Singaporean (reference level: G/G as 1; A-carriers as 2) samples, but three levels were considered for the analysis on the total sample (reference level: A/A as 1; A/G as 2; G/G as 3). The factor culture included two levels for the analysis developed on the total sample (reference level: Singaporean as 1; Italian as 2).

Overall, the hierarchical regression allowed identifying the positive predictors of the dependent variable at different stages of increasing complexity. The final step was reached when all the variables defined by the hypotheses were included in the model. As described, the final models included the interaction between all the embedded variables (interactive model) as predicted by the hypotheses.

This type of regression is particularly targeted in research design with multiple variables (e.g., different contributing factors and/or continuous subscales), nested designs, and unpredictable outcomes or exploratory analyses [10, 20] and was already applied in previous studies in this field [37, 38, 64, 63]. Before conducting the HMR analyses, the related assumptions (multicollinearity, i.e., variance inflation factor, homogeneity of variance, i.e., Bartlett’s test) were tested and satisfied. The post hoc statistical power was calculated for the final model of each series of HMR analysis of each sample using the G*Power software (Version 3.1) [21]. The magnitude of each effect for the hierarchical linear models was estimated by *R, R*^2^ and Δ*R*^2^.

For each sample, the level of significance for the effects emerging from the final step of each HMR analysis was adjusted as a function of the two HMR analyses conducted on anxiety and avoidance, respectively (2 repeated measures for the Italian, Singaporean, and total sample separately; corrected *alpha* = 0.025). Bonferroni’s method was chosen instead of alternative techniques to apply a more conservative correction.

For each sample, the significant main and interaction effects obtained from the final step of each series of HMR analysis were graphically represented by scatterplots with linear models and bar plots. A further assessment was advanced to inspect the direction of such effects. Pearson’s *r* and Fisher’s *z* tests evaluated the degree of association between the continuous independent variables and the dependent variables [16]. Variables related to PBI subscales were also divided into two groups (low vs. high PBI dimension) using the median split procedure for post hoc tests. The median values used to divide the PBI variables into the “high” and “low” subgroups are reported in Table 2. Post hoc analyses were computed for significant results on ECR-R variables adopting Student’s *t*-tests. Welch’s *t*-tests were preferred over Student’s *t*-tests when the variance of the comparing two groups from the normal population was different. A false discovery rate correction (e.g., Bonferroni’s method) for the *t*-tests was considered to circumscribe the probability of committing a type I error. The magnitude of the significant effects for the *t*-test was estimated by Cohen’s *d*.

With regard to the significant main effects, the mean differences of the given ECR-R variable between low and high PBI subscales (*α* = 0.05) were tested (e.g., see below the main effects for the Italian, Singaporean, and total sample). Concerning the significant interactions between genotype and environment, low vs. high PBI levels were compared to analyze the differences between the genetic carriers on the ECR-R dimension (e.g., see below the effects for the Singaporean sample given the corrected *α* = 0.025 and the total sample given the corrected *α* = 0.017). In the context of the significant interactions between culture and environment, low vs. high PBI levels were contrasted to probe the differences between the Italian and Singaporean individuals (corrected *α* = 0.025) on the ECR-R subscale (e.g., see below the effects for the total sample). In reference to the significant interactions between culture, genotype, and environment, low vs. high PBI levels were contrasted to inspect the differences between the genetic carriers from the Italian and Singaporean samples (corrected *α* = 0.025) on the ECR-R variable (e.g., see below the effects for the total sample).

## 3. Results

### 3.1. Preliminary Results

The Cronbach’s alpha coefficients ranged from 0.78 to 0.94, suggesting an overall very good internal consistency. In Table 2, all the Cronbach alpha values are listed, together with the main descriptive statistics of each sample. The psychometric features of the two instruments are compatible with those reported in a recent review on adult attachment assessment tools [57].

As observed in Table 2, acceptable skewness and kurtosis values, as well as visual inspection via density and quantile-quantile plots proved that the PBI and ECR-R subscales were normal univariate distributions (see Table 2). Only the distribution of PBI maternal care skewed right in the Italian sample (skewness = 1.15; kurtosis = 0.17); thus, the variable was square-transformed (skewness = −0.43; kurtosis = −0.88) before analysis. The same approach was adopted by previous research in the field [37].

A differential Bonferroni correction was applied to each group in adjusting *alpha* (Italian sample: 4 repeated measures, corrected *alpha* = 0.0125; Singaporean sample: 4 repeated measures, corrected *alpha* = 0.0125; total sample: 6 repeated measures, corrected *alpha* = 0.008). No effects of sex nor age on the ECR-R dimensions were significant (see Table 3). However, a main effect of culture was observed on adult anxiety, whereas the effect of culture on adult avoidance did not survive the correction (see Table 3; see Figure S3 of the Supplementary Materials). This preliminary result confirmed the necessity to inspect the potential interaction between culture, genotype, and caregiving behavior, as hypothesized by *HP3*.

**Table 3:**
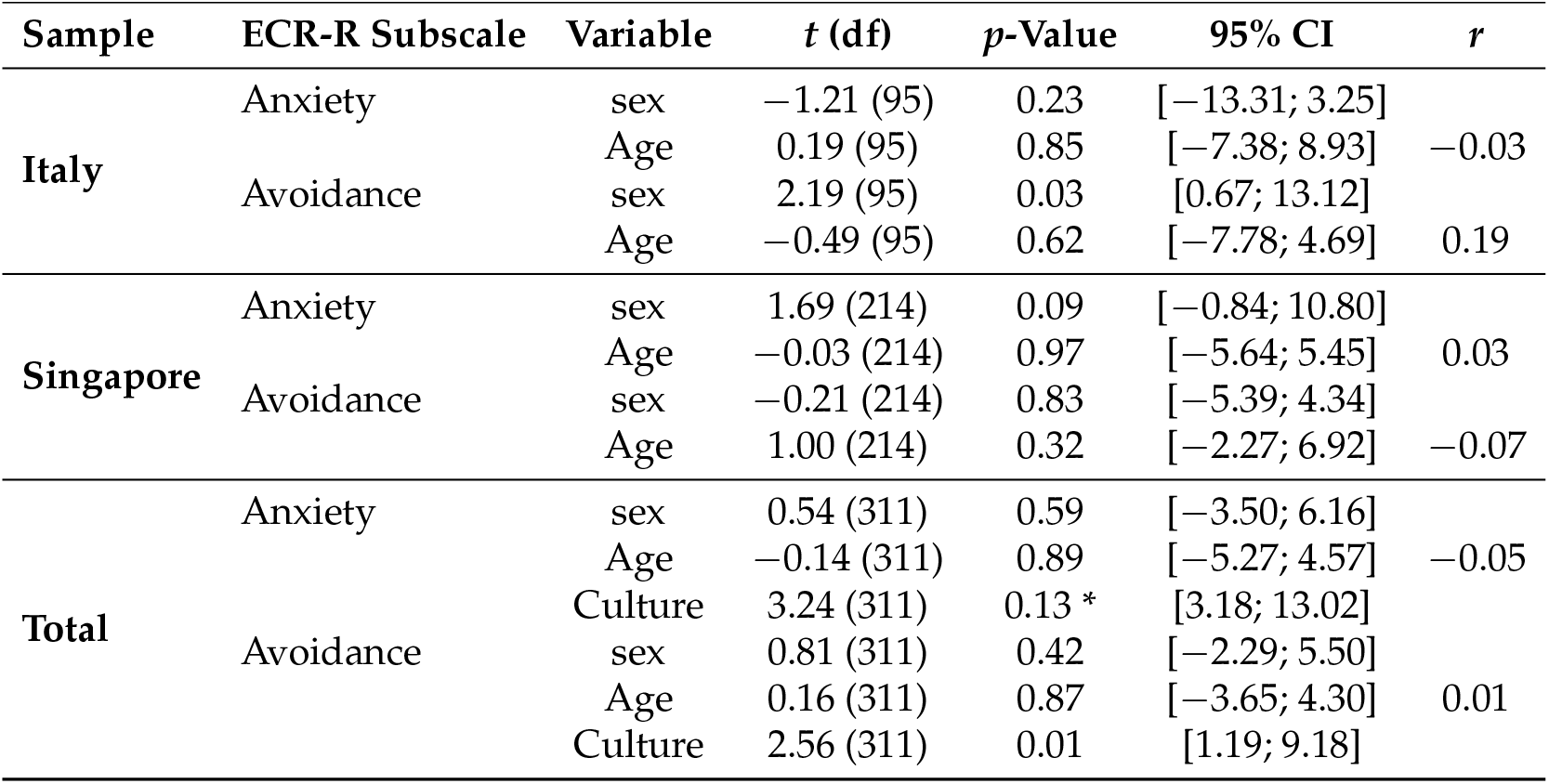
Summary of statistical values from Student’s *t*-tests for sex and age over the ECR-R dependent variables anxiety and avoidance. Pearson’s correlation coefficients among each ECR-R variable and age are also reported. * *p* < 0.008.

The means and standard deviations of the continuous variables are shown in Table 2, and their intercorrelations were examined, as reported in Table 4, for the Italian and Singaporean sample, respectively, and in Table S3 of the Supplementary Materials for the total sample (see Supplementary Materials).

**Table 4:** Pearson’s correlation coefficients and significance values among questionnaire subscales of the Italian and Singaporean samples separately. Values above the diagonal refer to the Singaporean sample, while values under the diagonal refer to the Italian sample. As regards the Italian sample, the PBI maternal care was square-transformed before the analysis. Significance is adjusted for multiple tests (corrected *alpha* = 0.003). * *p* < 0.003, ** *p* < 0.001, *** *p* < 0.0001.

| Subscale | M_Care | M_Overp | F_Care | F_Overp | Anxiety | Avoidance |
| --- | --- | --- | --- | --- | --- | --- |
| **M_Care** |  | −0.29 *** | 0.43 *** | −0.10 | −0.06 | −0.07 |
| **M_Overp** | −0.31 * |  | −0.17 | 0.39 *** | 0.1 | 0.05 |
| **F_Care** | 0.49 *** | −0.23 |  | −0.18 | −0.16 | −0.09 |
| **F_Overp** | −0.07 | 0.39 *** | −0.30 * |  | 0.17 | 0.10 |
| **Anxiety** | −0.36 ** | 0.42 *** | −0.37 ** | 0.20 |  | 0.38 *** |
| **Avoidance** | −0.25 | 0.19 | −0.20 | 0.08 | 0.35 ** |  |

A six-step HMR was conducted separately on the Italian and Singaporean sample for each ECR-R subscale (namely anxiety and avoidance) as the DV (see Table S2 of the Supplementary Materials). A seven-step HMR was conducted on the total sample for each ECR-R subscale as the DV (see Table S2 of the Supplementary Materials).

HMRs with the final step showing significant effects (*p* < 0.025) are explained below and reported in Table 5 for the Italian sample, in Table 6 for the Singaporean sample, and in Table 7 for the total sample. Otherwise, the explanation and tabulation of the results obtained from the HMRs not presenting significant effects (*p* > 0.025) at the final step are presented in the Supplementary Materials (see Tables S4–S6 of the Supplementary Materials).

**Table 5:** Hierarchical multiple regression on ECR-R anxiety for the Italian sample. Note. SE = standard error of the unstandardized coefficient. * *p* < 0.05; ** *p* < 0.01; *** *p* < 0.001.

| Step | Variable | $\beta$ | SE | $t$ | $R$ | $R^2$ | $\Delta R^2$ |
| --- | --- | --- | --- | --- | --- | --- | --- |
| 1 |  |  |  |  | 0.07 | 0.01 | 0.01 |
|  | OXTR rs53576 | 2.84 | 4.09 | 0.69 |  |  |  |
| 2 |  |  |  |  | 0.37 | 0.14 | 0.13 *** |
|  | OXTR rs53576 | 3.11 | 3.83 | 0.81 |  |  |  |
|  | M_Care | -0.02 *** | 0.01 | -3.81 |  |  |  |
| 3 |  |  |  |  | 0.49 | 0.24 | 0.10 *** |
|  | OXTR rs53576 | 3.36 | 3.62 | 0.93 |  |  |  |
|  | M_Care | -0.01 ** | 0.01 | -2.73 |  |  |  |
|  | M_OverP | 1.08 *** | 0.30 | 3.54 |  |  |  |
| 4 |  |  |  |  | 0.53 | 0.28 | 0.04 * |
|  | OXTR rs53576 | 4.06 | 3.55 | 1.14 |  |  |  |
|  | M_Care | -0.01 | 0.01 | -1.46 |  |  |  |
|  | M_OverP | 1.01 ** | 0.30 | 3.39 |  |  |  |
|  | F_Care | -0.50 * | 0.22 | -2.25 |  |  |  |
| 5 |  |  |  |  | 0.53 | 0.28 | 0 |
|  | OXTR rs53576 | 4.06 | 3.57 | 1.14 |  |  |  |
|  | M_Care | -0.01 | 0.01 | -1.42 |  |  |  |
|  | M_OverP | 1.01 ** | 0.33 | 3.11 |  |  |  |
|  | F_Care | -0.50 * | 0.23 | -2.13 |  |  |  |
|  | F_OverP | 0.00 | 0.29 | 0.00 |  |  |  |
| 6 |  |  |  |  | 0.54 | 0.29 | 0.01 |
|  | OXTR rs53576 | 18.08 | 17.73 | 1.02 |  |  |  |
|  | M_Care | -0.01 | 0.01 | -0.72 |  |  |  |
|  | M_OverP | 1.12 * | 0.47 | 2.40 |  |  |  |
|  | F_Care | -0.48 | 0.38 | -1.29 |  |  |  |
|  | F_OverP | 0.20 | 0.44 | 0.45 |  |  |  |
|  | OXTR rs53576 x M_Care | 0.00 | 0.01 | -0.34 |  |  |  |
|  | OXTR rs53576 x M_OverP | -0.26 | 0.67 | -0.39 |  |  |  |
|  | OXTR rs53576 x F_Care | -0.01 | 0.49 | -0.01 |  |  |  |
|  | OXTR rs53576 x F_OverP | -0.46 | 0.61 | 0.76 |  |  |  |

**Table 6:**
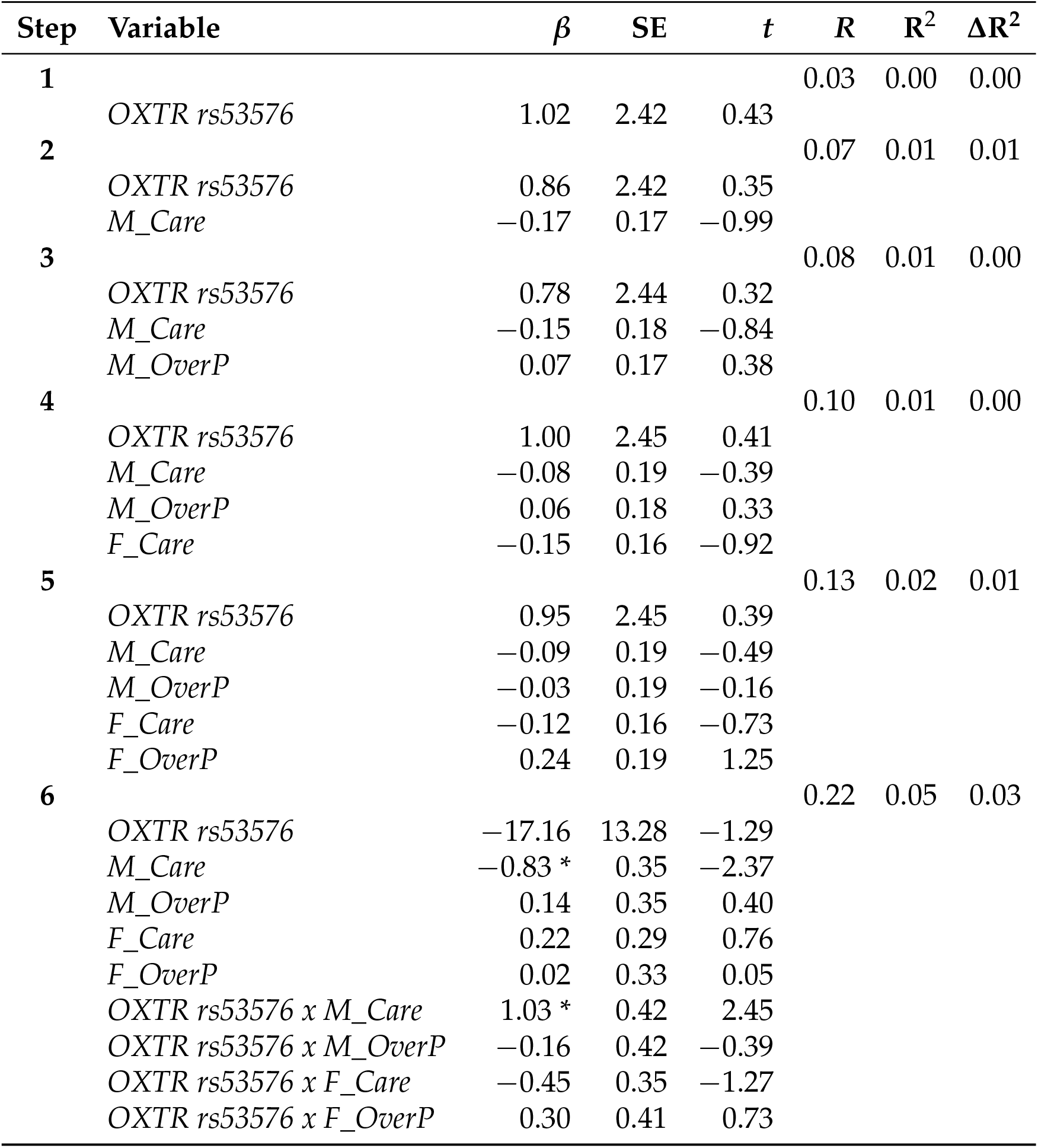
Hierarchical multiple regression on ECR-R avoidance for the Singaporean sample. Note. SE = standard error of the unstandardized coefficient. * *p* < 0.05.

| Step | Variable | $\beta$ | SE | $t$ | R | R<sup>2</sup> | $\Delta R^2$ |
| --- | --- | --- | --- | --- | --- | --- | --- |
| 1 |  |  |  |  | 0.03 | 0.00 | 0.00 |
|  | *OXTR rs53576* | 1.02 | 2.42 | 0.43 |  |  |  |
| 2 |  |  |  |  | 0.07 | 0.01 | 0.01 |
|  | *OXTR rs53576* | 0.86 | 2.42 | 0.35 |  |  |  |
|  | *M_Care* | -0.17 | 0.17 | -0.99 |  |  |  |
| 3 |  |  |  |  | 0.08 | 0.01 | 0.00 |
|  | *OXTR rs53576* | 0.78 | 2.44 | 0.32 |  |  |  |
|  | *M_Care* | -0.15 | 0.18 | -0.84 |  |  |  |
|  | *M_OverP* | 0.07 | 0.17 | 0.38 |  |  |  |
| 4 |  |  |  |  | 0.10 | 0.01 | 0.00 |
|  | *OXTR rs53576* | 1.00 | 2.45 | 0.41 |  |  |  |
|  | *M_Care* | -0.08 | 0.19 | -0.39 |  |  |  |
|  | *M_OverP* | 0.06 | 0.18 | 0.33 |  |  |  |
|  | *F_Care* | -0.15 | 0.16 | -0.92 |  |  |  |
| 5 |  |  |  |  | 0.13 | 0.02 | 0.01 |
|  | *OXTR rs53576* | 0.95 | 2.45 | 0.39 |  |  |  |
|  | *M_Care* | -0.09 | 0.19 | -0.49 |  |  |  |
|  | *M_OverP* | -0.03 | 0.19 | -0.16 |  |  |  |
|  | *F_Care* | -0.12 | 0.16 | -0.73 |  |  |  |
|  | *F_OverP* | 0.24 | 0.19 | 1.25 |  |  |  |
| 6 |  |  |  |  | 0.22 | 0.05 | 0.03 |
|  | *OXTR rs53576* | -17.16 | 13.28 | -1.29 |  |  |  |
|  | *M_Care* | -0.83 * | 0.35 | -2.37 |  |  |  |
|  | *M_OverP* | 0.14 | 0.35 | 0.40 |  |  |  |
|  | *F_Care* | 0.22 | 0.29 | 0.76 |  |  |  |
|  | *F_OverP* | 0.02 | 0.33 | 0.05 |  |  |  |
|  | *OXTR rs53576 x M_Care* | 1.03 * | 0.42 | 2.45 |  |  |  |
|  | *OXTR rs53576 x M_OverP* | -0.16 | 0.42 | -0.39 |  |  |  |
|  | *OXTR rs53576 x F_Care* | -0.45 | 0.35 | -1.27 |  |  |  |
|  | *OXTR rs53576 x F_OverP* | 0.30 | 0.41 | 0.73 |  |  |  |

**Table 7:**
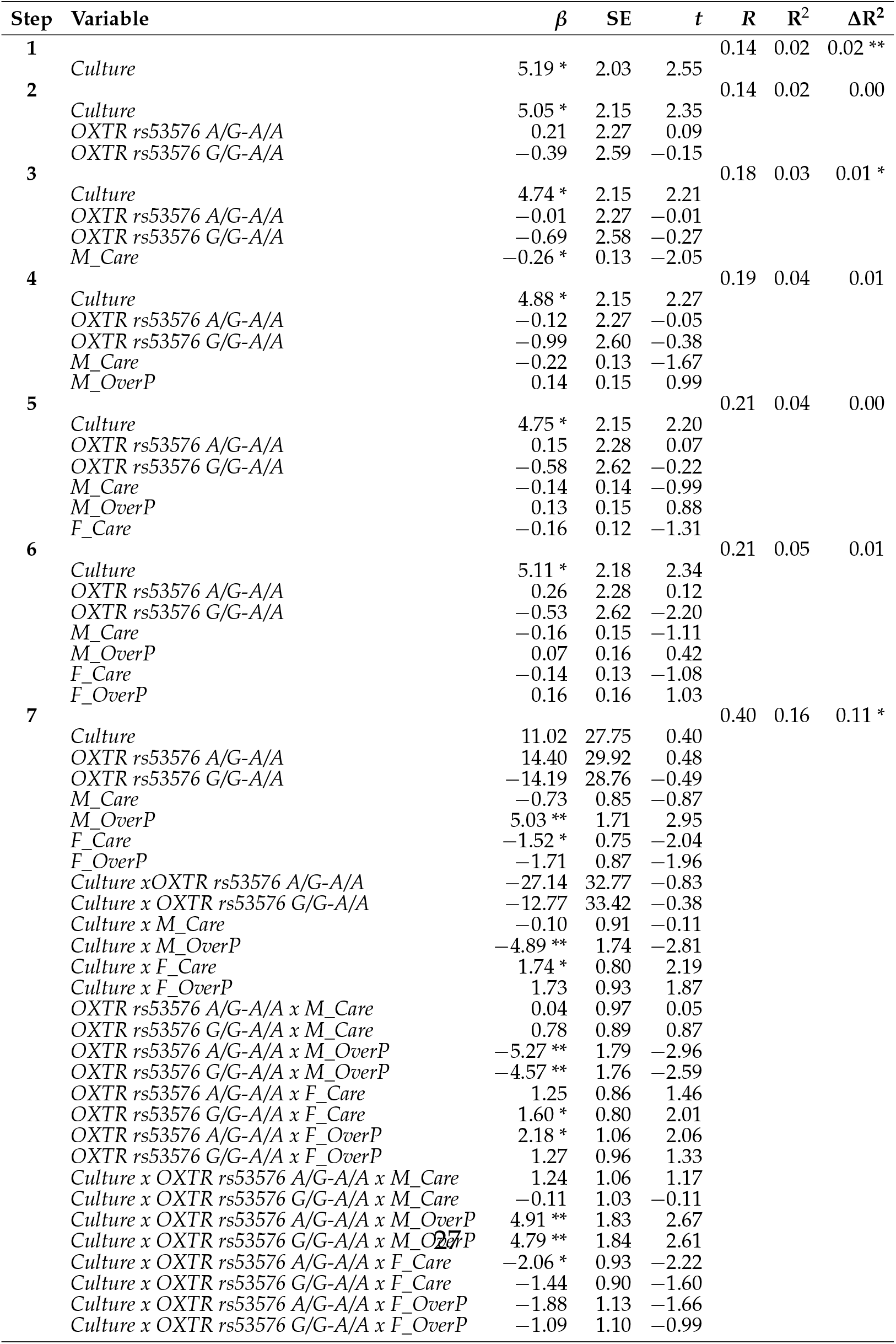
Hierarchical multiple regression on ECR-R avoidance for the total sample. Note. SE = standard error of the unstandardized coefficient. * *p* < 0.05, ** *p* < 0.01.

| Step | Variable | $\beta$ | SE | $t$ | $R$ | $R^2$ | $\Delta R^2$ |
| --- | --- | --- | --- | --- | --- | --- | --- |
| 1 |  |  |  |  | 0.14 | 0.02 | 0.02 ** |
|  | Culture | 5.19 * | 2.03 | 2.55 |  |  |  |
| 2 |  |  |  |  | 0.14 | 0.02 | 0.00 |
|  | Culture | 5.05 * | 2.15 | 2.35 |  |  |  |
|  | OXTR rs53576 A/G-A/A | 0.21 | 2.27 | 0.09 |  |  |  |
|  | OXTR rs53576 G/G-A/A | -0.39 | 2.59 | -0.15 |  |  |  |
| 3 |  |  |  |  | 0.18 | 0.03 | 0.01 * |
|  | Culture | 4.74 * | 2.15 | 2.21 |  |  |  |
|  | OXTR rs53576 A/G-A/A | -0.01 | 2.27 | -0.01 |  |  |  |
|  | OXTR rs53576 G/G-A/A | -0.69 | 2.58 | -0.27 |  |  |  |
|  | M_Care | -0.26 * | 0.13 | -2.05 |  |  |  |
| 4 |  |  |  |  | 0.19 | 0.04 | 0.01 |
|  | Culture | 4.88 * | 2.15 | 2.27 |  |  |  |
|  | OXTR rs53576 A/G-A/A | -0.12 | 2.27 | -0.05 |  |  |  |
|  | OXTR rs53576 G/G-A/A | -0.99 | 2.60 | -0.38 |  |  |  |
|  | M_Care | -0.22 | 0.13 | -1.67 |  |  |  |
|  | M_OverP | 0.14 | 0.15 | 0.99 |  |  |  |
| 5 |  |  |  |  | 0.21 | 0.04 | 0.00 |
|  | Culture | 4.75 * | 2.15 | 2.20 |  |  |  |
|  | OXTR rs53576 A/G-A/A | 0.15 | 2.28 | 0.07 |  |  |  |
|  | OXTR rs53576 G/G-A/A | -0.58 | 2.62 | -0.22 |  |  |  |
|  | M_Care | -0.14 | 0.14 | -0.99 |  |  |  |
|  | M_OverP | 0.13 | 0.15 | 0.88 |  |  |  |
|  | F_Care | -0.16 | 0.12 | -1.31 |  |  |  |
| 6 |  |  |  |  | 0.21 | 0.05 | 0.01 |
|  | Culture | 5.11 * | 2.18 | 2.34 |  |  |  |
|  | OXTR rs53576 A/G-A/A | 0.26 | 2.28 | 0.12 |  |  |  |
|  | OXTR rs53576 G/G-A/A | -0.53 | 2.62 | -0.20 |  |  |  |
|  | M_Care | -0.16 | 0.15 | -1.11 |  |  |  |
|  | M_OverP | 0.07 | 0.16 | 0.42 |  |  |  |
|  | F_Care | -0.14 | 0.13 | -1.08 |  |  |  |
|  | F_OverP | 0.16 | 0.16 | 1.03 |  |  |  |
| 7 |  |  |  |  | 0.40 | 0.16 | 0.11 * |
|  | Culture | 11.02 | 27.75 | 0.40 |  |  |  |
|  | OXTR rs53576 A/G-A/A | 14.40 | 29.92 | 0.48 |  |  |  |
|  | OXTR rs53576 G/G-A/A | -14.19 | 28.76 | -0.49 |  |  |  |
|  | M_Care | -0.73 | 0.85 | -0.87 |  |  |  |
|  | M_OverP | 5.03 ** | 1.71 | 2.95 |  |  |  |
|  | F_Care | -1.52 * | 0.75 | -2.04 |  |  |  |
|  | F_OverP | -1.71 | 0.87 | -1.96 |  |  |  |
|  | Culture x OXTR rs53576 A/G-A/A | -27.14 | 32.77 | -0.83 |  |  |  |
|  | Culture x OXTR rs53576 G/G-A/A | -12.77 | 33.42 | -0.38 |  |  |  |
|  | Culture x M_Care | -0.10 | 0.91 | -0.11 |  |  |  |
|  | Culture x M_OverP | -4.89 ** | 1.74 | -2.81 |  |  |  |
|  | Culture x F_Care | 1.74 * | 0.80 | 2.19 |  |  |  |
|  | Culture x F_OverP | 1.73 | 0.93 | 1.87 |  |  |  |
|  | OXTR rs53576 A/G-A/A x M_Care | 0.04 | 0.97 | 0.05 |  |  |  |
|  | OXTR rs53576 G/G-A/A x M_Care | 0.78 | 0.89 | 0.87 |  |  |  |
|  | OXTR rs53576 A/G-A/A x M_OverP | -5.27 ** | 1.79 | -2.96 |  |  |  |
|  | OXTR rs53576 G/G-A/A x M_OverP | -4.57 ** | 1.76 | -2.59 |  |  |  |
|  | OXTR rs53576 A/G-A/A x F_Care | 1.25 | 0.86 | 1.46 |  |  |  |
|  | OXTR rs53576 G/G-A/A x F_Care | 1.60 * | 0.80 | 2.01 |  |  |  |
|  | OXTR rs53576 A/G-A/A x F_OverP | 2.18 * | 1.06 | 2.06 |  |  |  |
|  | OXTR rs53576 G/G-A/A x F_OverP | 1.27 | 0.96 | 1.33 |  |  |  |
|  | Culture x OXTR rs53576 A/G-A/A x M_Care | 1.24 | 1.06 | 1.17 |  |  |  |
|  | Culture x OXTR rs53576 G/G-A/A x M_Care | -0.11 | 1.03 | -0.11 |  |  |  |
|  | Culture x OXTR rs53576 A/G-A/A x M_OverP | 4.91 ** | 1.83 | 2.67 |  |  |  |
|  | Culture x OXTR rs53576 G/G-A/A x M_OverP | 4.79 ** | 1.84 | 2.61 |  |  |  |
|  | Culture x OXTR rs53576 A/G-A/A x F_Care | -2.06 * | 0.93 | -2.22 |  |  |  |
|  | Culture x OXTR rs53576 G/G-A/A x F_Care | -1.44 | 0.90 | -1.60 |  |  |  |
|  | Culture x OXTR rs53576 A/G-A/A x F_OverP | -1.88 | 1.13 | -1.66 |  |  |  |
|  | Culture x OXTR rs53576 G/G-A/A x F_OverP | -1.09 | 1.10 | -0.99 |  |  |  |

### 3.2. Italian Sample

#### 3.2.1. Anxiety

Table 5 represents the series of HMRs conducted on the ECR-R anxiety of the Italian sample (total sum of squares - SS = 38,793). At Step 1, *OXTR* explained a non-significant 1% of variance (*F*(1,95) = 0.62, residual sum of squares - RSS = 38,597, *p* = 0.43). Entering the maternal care variable at Step 2 explained a significant additional 13.3% of variance (*F*(1,94) = 16.35, RSS = 33,425, *p* < 0.0001). After introducing maternal overprotection at Step 3, both maternal care and maternal overprotection were positive predictors, accounting for a significant 10% of variance (*F*(1,93) = 12.55, RSS = 29,455, *p* < 0.001). The addition of the paternal care variable to the regression model (Step 4) justified a further significant 4% of variation in anxiety (*F*(1,92) = 4.85, RSS = 27,921, *p* = 0.03) given the main effects of maternal overprotection and maternal care. No significant change in the R^2^ was observed when the final variable paternal overprotection was included at Step 5 (*F*(1,91) = 0.00, RSS = 27,921, *p* = 0.99). At Step 6, the hypothesized interaction between *OXTR* and paternal bonding dimensions on anxiety was tested (f^2^ = 0.39, power = 0.99). Here, a main effect of maternal overprotection was found (*β* = 1.11, SE = 0.46, *t* = 2.40, *p* < 0.02). Although the final model computed the highest proportion of explained variance in anxiety by the model (R^2^ = 0.29), no significant difference was detected compared to the previous step (*F*(1,91) = 0.32, RSS = 27,517, *p* = 0.87).

As indicated in Table 4, maternal overprotection was also found positively associated with adult anxiety (*t* = 4.46, df = 95, *r* = 0.42, *p* < 0.0001) (Figure 1). Specifically, the two-tailed post hoc Welch’s *t* test (*α* = 0.05) revealed that felt anxiety towards the partner was significantly different between participants who experienced low levels in maternal overprotection compared to those who had a past of maternal overprotection (*t* = −2.31, df = 71.3, *p* = 0.024, *d*= −0.49) (Figure 1). Means and standard error values for this effect on avoidance are available from Table S7 of the Supplementary Materials (see Supplementary Materials).

**Figure 1:**
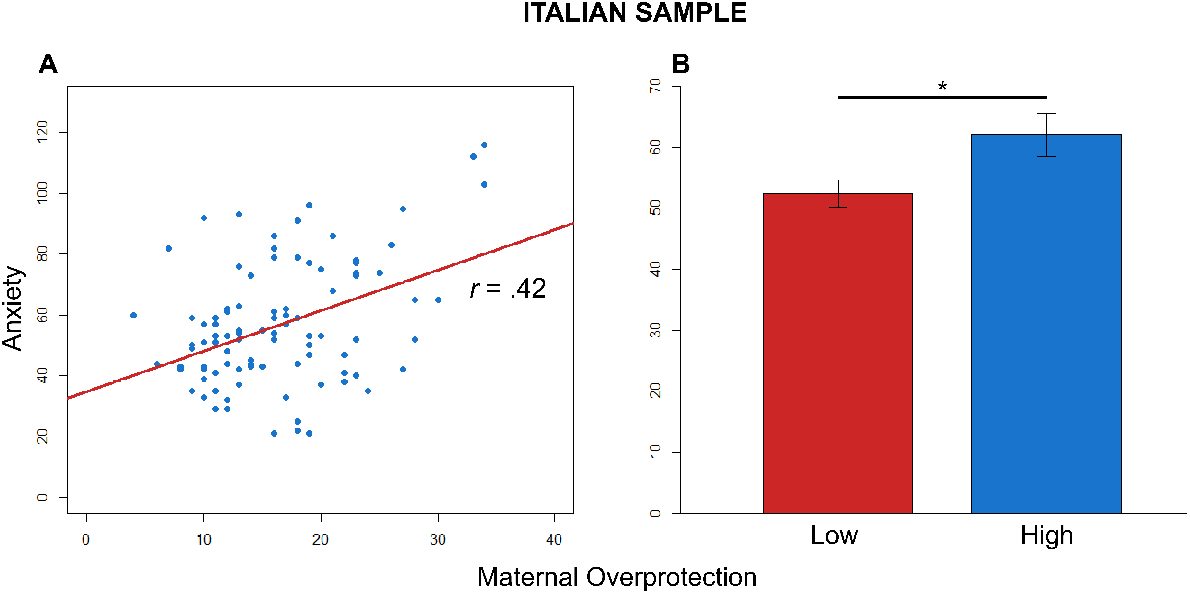
Effect on anxiety for the Italian sample. (**A**) Effect of early maternal overprotection on adult anxiety. Correlation between the scores of anxiety and maternal overprotection. Blue circles = Italian participants. The red line represents the linear model for the Italian participants. The *r*-value refers to Pearson’s *r* correlation. (**B**) Comparison between the scores of ECR-R anxiety in low and high maternal overprotection (* *p* < 0.025).

Contrary to *HP1*, no main effect of genotype or interactions between covariates and genotype were identified.

### 3.2.2. Avoidance

Table S4 of the Supplementary Materials (see Supplementary Materials) illustrates the series of HMRs conducted on ECR-R avoidance for the Italian sample. The related description is reported in the Supplementary Results (see Supplementary Materials).

### 3.3. Singaporean Sample

#### 3.3.1. Anxiety

Table S5 of the Supplementary Materials (see Supplementary Materials) shows the series of HMR conducted on ECR-R anxiety for the Singaporean sample. The related details are presented in the Supplementary Results (see Supplementary Materials).

### 3.3.2. Avoidance

Table 6 presents the series of HMRs performed on ECR-R avoidance for the Singaporean sample (total SS = 63,074). Entering at Step 1 the only genetic variable did not increase the explained portion of variance (*F*(1,214) = 0.18, RSS = 63,021, *p* = 0.67). At Step 2, maternal care was added to the regression model and accounted for a non-significant 1% of additional variance (*F*(1,213) = 0.99, RSS = 62,733, *p* = 0.32). The inclusion of maternal overprotection at Step 3 did not further improve the value of **R**^2^ (*F*(1,212) = 0.15, RSS = 62,689, *p* = 0.70). No change in the explained variance was even observed when introducing the paternal care dimension at Step 4 (*F*(1,211) = 0.87, RSS = 62,438, *p* = 0.35). At Step 5, the addition of the subscale paternal overprotection justified an increased 1% of variation in avoidance, which was not significant (*F*(1,210) = 1.59, RSS = 61,976, *p* = 0.21). At Step 6, the hypothesized (*HP2*) interaction between *OXTR* and paternal bonding dimensions on avoidance was tested (f^2^ = 0.05, power = 0.67). The final model maximized the explained variance (R^2^ = 0.05), but no significant difference was found compared to the previous step (*F*(4,206) = 1.74, RSS = 59,956, *p* = 0.14).

At this final step, early maternal care was the strongest predictor of adult avoidance (*β* = −0.83, SE = 0.35, *t* = −2.37, *p* < 0.02). As reported in Table 4, the negative correlation between maternal care and adult avoidance was not significant (*t* = −1.02, df = 214, *r* = −0.07, *p* = 0.30) (Figure 2). Specifically, the two-tailed post hoc Student’s *t* test (*α* = 0.05) showed that avoidance towards the partner was not significantly different between participants who reported low and high scores in maternal care (*t* = 0.78, df = 214, *p* = 0.44, *d*= 0.11) (Figure 2).

**Figure 2:**
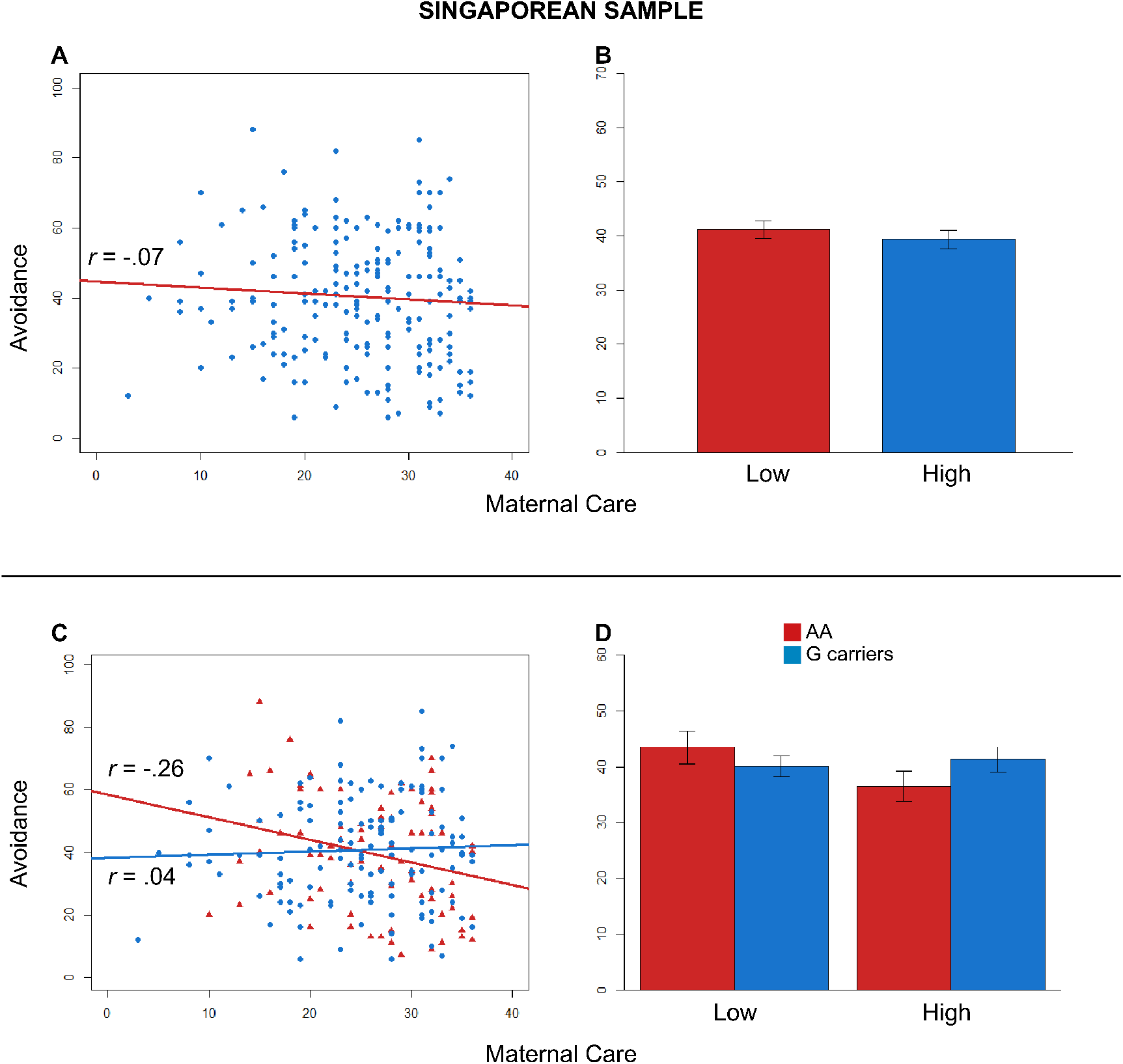
Effects on avoidance for the Singaporean sample. (**A**) Effect of early maternal care on adult avoidance. Correlation between the reported avoidance and maternal care. Blue circles = Singaporean participants. The red line represents the linear model for the Singaporean participants. (**B**) Comparison between the scores of ECR-R avoidance in low and high maternal care. (**C**) Effect of the interaction between genotype and early maternal care on adult avoidance. Correlations between the scores of anxiety and the recalled maternal care. Red triangles = A/A; blue circles = G-carriers. Lines depict the linear models for A/A homozygotes (red) and G-carriers (blue). (**D**) Contrast between the scores of ECR-R avoidance in A/A homozygotes (red) and G-carriers (blue) divided into low and high maternal care. *r*-values refer to Pearson’s *r* correlations for both (**A**,**C**).

The significant two-way interaction between rs53576 and maternal over-protection also emerged for adult avoidance (*β* = −0.83, SE = 0.35, *t* = −2.37, *p* < 0.02). The distribution of genotypes A/A vs. G-carriers was not significantly different between high vs. low maternal care (*X*^2^ (1) = 1.19, *p* = 0.028). Maternal care was positively associated with avoidance for G-carriers (*t* = 0.51, df = 134, *r* = 0.04, *p* = 0.61), but negatively associated with avoidance for A/A homozygotes (*t* = −2.38, df = 78, *r* = −0.26, *p* = 0.02), as predicted by *HP2* (Figure 2). Although only one Pearson’s *r* was significant, Fisher’s z test confirmed that the difference between the slopes for A/A and G-carriers was significant (*z* = −2.14, *p* = 0.03). The one-tailed post hoc Student’s *t* test (corrected *α* = 0.025) revealed that adult avoidance was not significantly different between A/A- and G-carriers when they experienced a history of low maternal caregiving (*t* = −1.01, df = 107, *p* = 0.84, *d*= −0.21) (Figure 2).

Means and standard error values for each effect on avoidance are available from Table S8 of the Supplementary Materials (see Supplementary Materials).

### 3.4. Total Sample: Italian and Singaporean Participants

#### 3.4.1. Anxiety

Table S6 of the Supplementary Materials (see Supplementary Materials) represents the series of HMRs conducted on ECR-R anxiety for the total sample. The statistical details are indicated in the Supplementary Results (see Supplementary Materials).

#### 3.4.2. Avoidance

Table 7 reports the series of HMRs performed on ECR-R avoidance for the total sample (total SS = 87,563). The regression revealed that at Stage 1, culture contributed significantly to the model explaining the first 2% of variance (*F*(1,311) = 6.90, RSS = 85,763, *p* = 0.009). Culture was significant at this step, as a positive predictor. At Step 2, introducing the genetic factor did not increase the variance of avoidance (*F*(2,309) = 0.03, RSS = 85,745, *p* = 0.97), but a main effect of culture was still present. At Step 3, maternal care was added to the regression model and accounted for a significant 1% of additional variance (*F*(1,308) = 4.42, RSS = 84,592, *p* = 0.04). Here, the variable culture and the novel variable maternal care were the strongest predictors. At Step 4, the improvement of 1% of variation in avoidance due to the inclusion of maternal overprotection in the model was not significant (*F*(1,307) = 1.03, RSS = 84,325, *p* = 0.31). Only culture was a positive predictor at this stage. No change in the explained variance was observed when introducing the paternal care dimension at Step 5 (*F*(1,306) = 1.79, RSS = 83,858, *p* = 0.18). Culture was still a strong predictor at this stage. The addition of the paternal overprotection variable to the regression model at Step 6 justified 1% of the variation in avoidance which did not result in being statistically significant *F*(1,305) = 1.11, RSS = 83,569, *p* = 0.29). However, a significant main effect of culture was identified at this stage. Finally, at Step 7, the potential interaction between culture, *OXTR*, and PBI dimensions on avoidance (*H3*) was inspected (f^2^ = 0.17, power = 0.99). This interactive model increased the explained variance (R^2^ = 0.16) significantly from the previous step (*F*(22,283) = 1.70, RSS = 73,827, *p* = 0.028).

At this last step, one one-way effect, three two-way interaction effects, and two three-way interaction effects were found. Means and standard error values for each effect on avoidance are available from Table S9 of the Supplementary Materials (see Supplementary Materials).

Firstly, early maternal overprotection singularly predicted adult avoidance (*β* = 5.03, SE = 1.71, *t* = 2.95, *p* < 0.004). As evident from Table S3 of the Supplementary Materials (see Supplementary Materials), the positive correlation between maternal overprotection and adult avoidance was not significant (*t* = −1.26, df = 311, *r* = 0.07, *p* = 0.21) (Figure 3A; blue circles = Italian and Singaporean participants merged; the red line represents the linear model for all the participants). The two-tailed post hoc Student’s *t* test (*α* = 0.05) showed that avoidance towards the partner was not significantly different between participants who reported low and high scores in maternal overprotection (*t* = −1.27, df = 311, *p* = 0.21, *d*= −0.14) (Figure 3B).

**Figure 3:**
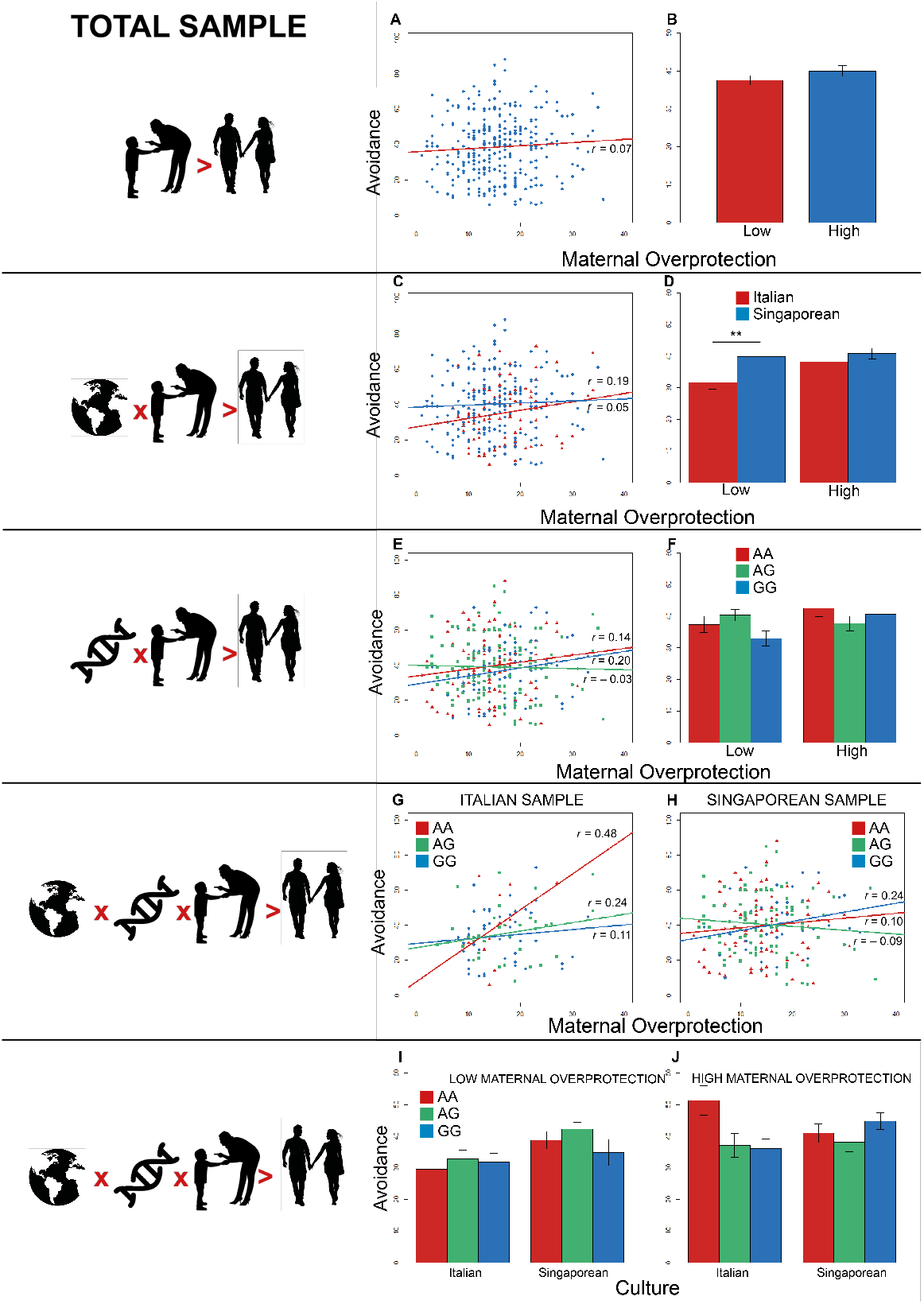
Effects on avoidance for the total sample. (**A**) Effect of early maternal overprotection on adult avoidance. (**B**) Comparison between the scores of ECR-R avoidance in low and high maternal overprotection. (**C**) Effect of the interaction between culture and early maternal overprotection on adult avoidance. (**D**) Contrast between the scores of ECR-R avoidance in Italian (red) and Singaporean (blue) groups. (**E**) Effect of the interaction between genotype and early maternal overprotection on adult avoidance. (**F**) Contrast between the scores of ECR-R avoidance in A/A homozygotes (red), A/G heterozygotes (green), and G/G homozygotes (blue) divided into low and high maternal overprotection. (**G**) Effect of the interaction between genotype and early maternal overprotection on adult avoidance for the Italian participants. (**H**) Effect of the interaction between genotype and early maternal overprotection on adult avoidance for the Singaporean participants. (**I**) Comparison between the scores of ECR-R avoidance in A/A homozygotes (red), A/G heterozygotes (green), and G/G homozygotes (blue) divided into Italian and Singaporean participants for low maternal overprotection. (**J**) Contrast between the scores of ECR-R avoidance in A/A homozygotes (red), A/G heterozygotes (green), and G/G homozygotes (blue) divided into Italian and Singaporean participants for high maternal overprotection. *r*-values refer to Pearson’s *r* correlations. (** *p* < 0.025).

The main effect of paternal care (*p* > 0.025) did not survive the magnitude of the Bonferroni correction (corrected *alpha* = 0.025).

As concerns the interaction effects, the significant two-way interaction between culture and maternal overprotection also emerged for avoidance (*β* = −4.89, SE = 1.74, *t* = −2.81, *p* = 0.005). Maternal overprotection positively correlated with the avoidance reported by both Italian (*t* = 1.91, df = 95, *r* = 0.19, *p* = 0.06) and Singaporean participants (*t* = 0.69, df = 214, *r* = 0.05, *p* = 0.49) (Figure 3C; red triangles = Italian participants; blue circles

= Singaporean participants; lines depict the linear models for Italian (red) and Singaporean (blue) participants). However, the slopes for Italians and Singaporeans were not significantly different (*z* = 1.15, *p* = 0.13). The exploratory two-tailed Student’s *t* test (corrected *alpha* = 0.025) revealed that adult avoidance was significantly different between Italian and Singaporean participants when they experienced a history of low maternal overprotection (*t* = −2.91, df = 162, *p* = 0.004, *d*=-0.51) (Figure 3D), but not when they had a past of high maternal overprotection (*t* = −0.88, df = 147, *p* = 0.38, *d*= −0.15).

The two-way interaction between culture and paternal care (*p* > 0.025) was not considered significant once *alpha* correction was applied to the linear models (corrected *alpha* = 0.025).

Two further significant two-way interactions between *OXTR* levels and *p* maternal overprotection on adult avoidance were detected (A/G-A/A: *β* = −5.27, SE = 1.71, *t* = −2.96, = 0.003; G/G-A/A: *β* = −4.57, SE = 1.76, *t* = −2.59, *p* = 0.009). The distribution of genotypes A/A vs. A/G vs. G/G was not significantly different between high vs. low maternal overprotection (*X*^2^ (2) = 3.75, *p* = 0.15). Maternal overprotection was positively associated with avoidance for A/A (*t* = 1.32, df = 92, *r* = 0.14, *p* = 0.19) and G/G homozygotes (*t* = 1.91, df = 88, *r* = 0.20, *p* = 0.06), but not for the heterozygotes (*t* = −0.31, df = 127, *r* = −0.03, *p* = 0.75) (Figure 3E; red triangles = A/A; green squares = A/G; blue circles = G/G; lines represent the linear models for A/A homozygotes (red), A/G (green), and G/G (blue)). No difference between the slopes for A/A and A/G (*z* = −.24, *p* = 0.21), for A/A and G/G (*z* = −0.44, *p* = 0.66), and for A/G and G/G (*z* = −1.67, *p* = 0.09) was observed. In the only condition of high maternal overprotection (corrected *α* = 0.017), the exploratory two-tailed post hoc Student’s *t* tests proved that adult avoidance was not significantly different between A/A and A/G (*t* = 1.34, df = 97, *p* = 0.18, *d*= 0.28), A/A and G/G (*t* = 0.58, df = 87, *p* = 57, *d*= 0.12), and A/G and G/G (*t* = 0.90, df = 108, *p* = 37, *d*= 0.17) (Figure 3F).

Considerably, the two two-way interaction respectively between *OXTR* and paternal care and between *OXTR* and paternal overprotection on adult avoidance did not fall below the level of acceptability (*p* > 0.025).

Concerning the potential three-way interaction expected for the observational *HP3*, two significant three-way interactions between culture, *OXTR* levels, and maternal overprotection on adult avoidance were discovered (A/G-A/A: *β* = 4.91, SE = 1.83, *t* = 2.67, *p* < 0.008; G/G-A/A: *β* = 4.79, SE = 1.84, *t* = 2.61, *p* = 0.009). In the Italian group, maternal overprotection was positively associated with avoidance for all the genetic groups (A/A: *t* = 1.86, df = 12, *r* = 0.48, *p* = 0.09; A/G: *t* = 1.39, df = 33, *r* = 0.24, *p* = 0.17; G/G: *t* = 0.73, df = 46, *r* = 0.11, *p* = 0.47) (Figure 3G; red triangles = A/A; green squares = A/G; blue circles = G/G; lines show the linear models for A/A homozygotes ((red), A/G (green), and G/G (blue)). No difference between the slopes for all the Italian genetic groups was found (A/A-A/G: *z* = 0.80, *p* = 0.21); A/A-G/G: *z* = 1.23, *p* = 0.11; A/G-G/G: *z* = 0.58, *p* = 0.28). In the Singaporean group, maternal overprotection was positively associated with avoidance for A/A (*t* = 0.90, df = 78, *r* = 0.10, *p* = 0.37) and G/G (*t* = 1.59, df = 40, *r* = 0.24, *p* = 0.12), but the same PBI dimension was negatively associated with avoidance for A/G (*t* = −0.85, df = 92, *r* = −0.09, *p* = 0.40) (Figure 3H; red triangles = A/A; green squares = A/G; blue circles = G/G; lines show the linear models for A/A homozygotes (red), A/G (green), and G/G (blue)). Fisher’s *z*-tests confirmed a significant difference between slopes for the A/G-G/G groups (*z* = 1.75, *p* = 0.04), but not for the A/A-A/G (*z* = 1.23, *p* = 0.11) and the A/A-G/G (*z* = −0.74, *p* = 0.23) groups. In the condition of low maternal overprotection, the two-tailed post hoc Student’s *t* tests (corrected *alpha* = 0.025) confirmed that adult avoidance was not significantly different between A/A and G/G both in the Italian (*t* = − 0.36, df = 30, *p* = 0.72, *d*=-0.15) and Singaporean group (*t* = 0.74, df = 61, *p* = 0.47, *d*= 0.21) (Figure 3I). Likewise, in the condition of high maternal overprotection (corrected *alpha* = 0.025), adult avoidance was not significantly different between A/A and G/G both in the Italian (*t* = 2.28, df = 28, *p* = 0.03, *d*= 1.04) and Singaporean group (*t* = 0.94, df = 57, *p* = 0.35, *d*=-0.25) (Figure 3J).

Nevertheless, the last three-way interaction between culture, *OXTR*, and paternal care approached, but did not reach significance (*p* > 0.025).

## 4. Discussion

Early interactions have been widely proven to have a crucial role in shaping psychological mechanisms that drive social interactions later in life. However, it is fundamental to take into account several factors that occur in the development of these psychosocial processes, such as genetic and cultural contributions. Starting from the evidence in the literature suggesting the key role of oxytocin [18, 19] and parental bonding in influencing social behavior [22, 14], the main purpose of this study was to examine the effects of the different genetic susceptibility and early interactions on the levels of anxiety and avoidance experienced in adult relationships in two different cultural contexts. Within this multicultural framework, the Italian and Singaporean samples were respectively considered as representative of a Western-dominant culture and an Eastern-dominant culture. Specifically, we focused on recalled parental bonding, assessed in terms of parental care and overprotection. According to the information on the mean and median of each subscale (see Table 2), while parental dimensions did not diverge between the two cultural groups, anxiety and avoidance experienced in adult relationships showed different means and median values in the two populations. ECR-R has been translated into and validated in other languages and used for investigations in Eastern countries, proving good internal validity. At the same time, the few studies that adopted ECR-R for cross-cultural comparisons [32, 44] highlighted the possibility that cultural differences could lay at the semantic level. Moreover, it is likely that participants belonging to the Singaporean sample, representing a collectivist population, could express preoccupation towards other people’s opinions compared to the Italian group, displaying greater levels of anxiety and avoidance [32]. However, due to the paucity of references regarding this point, results need to be interpreted cautiously. Additionally, we included in our analyses the effect of allelic variation of one *OXTR* gene SNP, namely *rs53576*. We were expecting to find effects for the interaction between the genetic and perceived caregiving variables, with different influences in the groups (tested in *HP1* and *HP2*). Moreover, we explored a more intertwined model, adding the influence of the cultural environment (tested in *HP3*).

### 4.1. Gene-By-Environment Interaction within the Western-Oriented Sample and the Eastern-Oriented Sample Evaluated Independently

With regards to the Italian sample, the first step considered only the *OXTR* variation in explaining the levels of anxiety and avoidance. Perceived maternal features were progressively entered into the model, both resulting as strong predictors for anxiety, with care displaying a negative association and overprotection a positive one. In the subsequent steps, recalled paternal characteristics were added to the model, and the care dimension for the father resulted as a predictor for anxiety as well. This might reflect the diversified, but equally important, contributions that both parents produce in their offspring, leading to different outcomes as a consequence [61]. In the model of interaction tested at the last stage, recalled maternal overprotection appeared to explain levels of anxiety, with a positive correlation: the greater the overprotection felt, the higher the levels of experienced anxiety. Here, Italian individuals with a past of perceived maternal overprotection reported higher levels of anxiety than those who recalled a less controlling relationship with their mothers. Perceived maternal control during childhood, assessed in terms of overprotection, seemed to be the construct that better explained the level of anxiety in adult relationships. This result was in line with the theoretical framework of the development of attachment [9], and furthermore, it was also supported by other studies [51, 61, 56, 35, 68]. The results suggested that both parents, to different extents, contributed in explaining the formation of anxiety-related patterns in adult social behavior. With regards to the experienced avoidance in adulthood, we found that maternal care contributed to the unique variance as a predictor, negatively associated. Although no other variables alone were proven robust predictors, the interaction between the genotype and early parental bonding dimensions substantially increased the explained variance in adult avoidance. However, due to the choice of a highly conservative correction method, the effect of maternal care emerging from the interactive model tested on avoidance did not reach the significance level. Interestingly, in the Italian sample, features of the mother appeared to influence the formation of psychological constructs of close relationships more than those of the father, highlighting the importance of the specific parental role in explaining outcomes in adult relationships [39, 28]. Moreover, within the final stages of the HMR, no significant effect was found for the interaction between *OXTR rs53576* and the parental bonding dimension on anxiety or avoidance, not confirming the first hypothesis.

Overall, these results suggest that the quality of familiar bonds in this population, where higher levels of expressed emotions often characterize the relationship between parents and offspring [53, 49, 40], could affect the experience of close relationships in adulthood more independently of the genetic sensitivity, invalidating the starting *HP1*.

Concerning the Singaporean sample, the development of HMRs on this population overall revealed distinct patterns of parental boding and genotype on adult attachment as a potential effect of Eastern cultures. As for adult anxiety, recalled paternal care and overprotection contributed more significantly to the model when respectively entered. Although the interaction between genotype and parental features did not improve the model in terms of the total amount of variance, an interaction effect between genotype and maternal overprotection emerged at this final stage. Such an effect was marginally significant, thus did not persist when the correction was applied.

Unlike the Italian sample, neither genotype, nor the recalled parental features resulted as a significant predictor of adult avoidance across the different steps of the model whose variance remained stable. In the final step, where all the targeted variables were included, maternal care was the strongest predictor of avoidance in adult close relationships with a negative trend. Moreover, the interaction between *OXTR* variations and maternal care reached statistical significance as well. In particular, the two variations followed a mutually opposite pattern of association, displaying a slight positive correlation for maternal care over avoidance levels for G-carriers and a negative one for A/A homozygotes. In line with *HP2*, the present cross-interaction supported the assumption of a differential sensitivity to environmental influences among carriers of distinctive genetic phenotype in *OXTR rs53576* [19, 18, 5]. While evidence suggested overall effects of *OXTR rs53576* on prosocial mechanisms, such as empathy or seeking emotional support, outcomes on attachment style were not consistent. Our results may reflect that the boundary condition plays a stronger role when it comes to adult attachment dimensions [30, 41, 59]; thus, caution is needed in addressing the A/A variant as a risk factor for non-optimal patterns of attachment. Currently, there are few studies examining the effects of the interaction of recalled parental bonding and *OXTR rs53576* on adult attachment dimensions in Eastern populations, considering the ethnic group separately [41, 59, 29], advocating the need for a focus on the importance of biological differences and distinct allelic distributions.

We observed that the effects in the two groups differed, especially in the relationship between recalled maternal bonding and the attachment dimension in adult relationships, reflecting the typical traits of familiar bonds: with regards to the Italian sample, where great attention is projected on the child and levels of emotions expressed are high [40], the construct of overprotection was associated with anxiety in adult relationships. According to the traits depicting individualistic cultures, the person is expected to be self-reliant and not to depend on other people for support, which in turn might be reflected in greater anxiety expressed or experienced, instead of avoidance [70]. On the other hand, in the Singaporean sample, where interdependence represents a typical trait in social groups, starting from the family, the avoidant pattern of behavior can describe a greater form of distress in relationships, compared to Western-oriented groups [70, 26].

### 4.2. Gene-By-Culture Interactions in the Total Sample

While the gene-by-environment framework focuses predominantly on external factors that characterize individual experience, the gene–culture interaction field of research focuses on the socio-cultural context. Within this theoretical frame, variants conferring genetic susceptibility can amplify typical cultural tendencies [37], such as interdependence [43]. Adopting HMRs, we aimed to explore the influence of the cultural environment and of its interaction with genetic variations sensitive to external context, such as perceived parental caregiving. In reference to the HMRs developed on adult anxiety, culture was the first variable added to the model, explaining a significant portion of variance. The subsequent addition of genotype did not increase the variation in anxiety, but this increase was true for the addition of maternal and paternal care, as well as paternal overprotection at different steps, respectively. Finally, the interaction among the culture, genotype, and parental bonding dimensions maximized the variance on adult anxiety. Despite this achievement, the model of interaction did no not reveal consistent effects on adult anxiety.

In the context of the HMRs advanced for adult avoidance, culture still was a key variable that was able to increase the variance significantly. Discordant with the significant effects of the numerous parental features on the variation in anxiety, only maternal care enlarged the amount of explained variation in avoidance. In accordance with the final model observed from the HMRs on anxiety, the highest quantity of total variance was reached when the interaction among the culture, genotype, and parental bonding dimensions was verified on avoidance.

From the model of the three-way interaction, multiple effects were observed. Firstly, maternal overprotection positively predicted adult avoidance when the cultural and genotype influences were not further examined. Secondly, the cultural component played a dominant role over experienced avoidance in adult close relationships. We observed that the effects on the two groups were different, especially in the relationship between culture and maternal overprotection. Specifically, maternal overprotection was positively associated with adult avoidance for both Western and Eastern samples. However, the Western-oriented participants reported lower mean levels of avoidance from the partner than the Eastern-oriented individuals. Mousavi and colleagues, although framing their exploration in a wider perception of anxiety, including social phobia, which is related to avoidant patterns of behavior, provided evidence that the perception of parental control or overprotection was reported in a stronger extent in a Caucasian sample [47].

Thirdly, the contributing action of the genetic predisposition driven by *OXTr rs53576* was also identified. In this regard, the significant interaction between genotype and maternal overprotection predicted adult avoidance when the participants were conceived together and independently from the cultural belonging to the group. However, the proper interpretation of this effect faced several difficulties in light of the different alleles’ frequency between Western and Eastern samples.

Analyses referring to the gene-by-culture frameworks displayed an interesting result: the different levels of associations between the genetic carriers A/A, A/G, and G/G and maternal overprotection on the adult avoidance for the Western-oriented and Eastern-oriented sample disclosed peculiar attachment patterns as a function of a differential genetic susceptibility to the external environment [36, 37]. Here, the less common variation (A/A for the Italian sample, G/G for the Singaporean group) was the one more directly correlated to greater levels of avoidance for higher levels of perceived maternal overprotection. Similar results were found by Kim and colleagues when exploring the role of the culture using a gene-by-environment approach between individualistic (American) and collectivist (Korean) populations in investigating emotion suppression [36]. Avoidance, likewise emotion suppression, can be described as a coping strategy to deal with stressful experiences [33]. Analogous to the findings of Kim and colleagues, greater levels of avoidance were observed among G/G-carriers in the Singaporean sample, compared to those carrying two copies of the A variant. Conversely, in the Italian group, G/G-carriers showed significantly lower levels of avoidance compared to the A/A-carriers. These findings suggested that the allelic variation retained to confer greater sensitivity to environmental input was correlated with more optimal outcomes involving social and emotional strategies, according to the specific societal context [37, 36]. Interestingly, in the Singaporean sample, people carrying the A/G variation displayed an opposite trend in terms of negative relationships with avoidance compared to both the positive associations of A/Aand G/G-carriers with the same ECR-R dimension. In the literature, evidence reports results for the heterozygous variation together with the less common homozygous pair of alleles in order to correct skewed distributions [36]; hence, this outcome requires caution when it comes to interpretation. Unexpectedly, the trend of avoidance scores in the Singaporean sample decreased for A/G-carriers, but increased for A/A and G/G homozygotes, suggesting a broad variability in perceived maternal control over close adult relationships.

## 5. Conclusions

In this study, we focused on two different social environments: the family, within witch the person experiences early relationships and builds the first behavioral patterns, and the cultural context, in which the grown person adopts a combined set of individual and shared criteria in relating to others. One of the strengths of this study was to provide a further step in comparing two distinct environments, each consisting of parenting strategies and family composition. Another important element considered was the distinction between maternal and paternal bonding. In fact, although currently, the competence regarding child care is becoming more and more equal between the two parents, fatherhood is an area that is still under-explored, especially in reference to the temporal dynamics of attachment in adulthood. Avoidance might have been more strongly correlated as it could play a mediating role between recalled parental features and anxiety [27]. In our approach, we divided the different combination of the single variants for the Italian and Singaporean group, in order to maximize the effects of the genetic distribution in the different ethnic groups. This study also presented limitations that are important to consider, starting from the cross-sectional approach that highlighted a weaker reliance on retrospective reporting of parental bonding, compared to a longitudinal one. Another point is that, although we utilized original (for Singaporean group) and validated translated (for Italian group) versions of the questionnaires, some conceptual understanding issues could have led to different interpretations of the items, hence to altered scoring that might have influenced the results. Additionally, the numerosity of the sample represents a challenge in gene-by-environment interaction studies. In fact, limited to the allelic distribution in the total sample, which was comprised of both the Italian and the Singaporean sample, the frequency of A/A and G/G homozygotes and A/G heterozygotes was unbalanced, and this may have influenced the outcomes. Finally, for both of the countries, the sample was composed by undergraduate/post-graduate students, mainly belonging to social science or psychological fields, who happened to be familiar with the theories beneath the study. Overall, while this study represents a further step in understanding the development of bonding mechanisms in different cultural contexts, it is at the same time a starting point for further exploration where more modern approaches could be combined, in order to have a more comprehensive understanding of the factors that might affect the dynamics of attachment, in both early stages and subsequent phases of life. Future research should include a larger number of participants in both populations, in order to better analyze the impact of the three genotypes and compare the results from the present study. Further investigations should extend the assessment by integrating psychological mechanisms underlying social behavior and expectations, which may play a mediating role between childhood reminiscence of parental care and experienced anxiety or avoidance in adulthood. Moreover, adult social relationships cannot exclude online social interactions; exploring the interaction between genes and environmental contexts over the experience of relationships in a virtual domain would allow researchers to have access to a more structured and complete view of human relationships [6, 13]. The implications of our findings for clinical work could cover an expanding area of interest in line with the current biopsychosocial model of health and psychological wellness, that is the influence of cultural components and early interactions in understanding the development of social behavior and related disorders. This would allow practitioners in child developmental and parenting fields to formulate interventions that take into account also the cultural features of the person, which is a relevant topic, due to the increasing mobility of single people and families.

## Supporting information

https://www.mdpi.com/2076-3425/11/4/496/s1

## 6. Supplementary Materials

Supplementary Results: Italian sample: avoidance; Singaporean sample: anxiety; total sample: anxiety. Supplementary Tables: Table S1: Age and sex differences between the rs53576 genetic groups for the Italian, Singaporean, and total sample. Table S2: Models applied to the six-step hierarchical multiple regressions on the Italian, Singaporean, and total sample; Table S3: Correlation matrix among questionnaire subscales across all participants; Table S4: Hierarchical multiple regression on adult avoidance for the Italian participants; Table S5: Hierarchical multiple regression on adult anxiety for the Singaporean participants; Table S6: Hierarchical multiple regression on adult anxiety for all the participants; Table S7: Means and standard errors of ECR-R anxiety by the independent variables for the Italian participants; Table S8: Means and standard errors of ECR-R avoidance by the independent variables for the Singaporean participants; Table S9: Means and standard errors of ECR-R avoidance by the independent variables for all the participants assessed. Supplementary Figures: Figure S1: Comparison between levels of sex and age on anxiety and avoidance for the Italian participants; Figure S2: Comparison between levels of sex and age on anxiety and avoidance for the Singaporean participants; Figure S3: Comparison between levels of sex, age, and culture on anxiety and avoidance for all the participants assessed.

## 7. Authors contribution

Conceptualization, I.C., B.L. and G.E.; methodology, A.B., I.C., B.L. and G.E.; formal analysis, A.B. and J.N.F.; investigation, A.B., I.C., B.L. and G.E.; data curation, I.C.; writing–original draft preparation, I.C. and A.B.; writing–review and editing, I.C, A.B., B.L., P.S. and G.E.; visualization, A.B.; supervision, B.L. and G.E.; funding acquisition, G.E. All authors have read and agreed to the published version of the manuscript.

## 9. Acknowledgements

The authors would like to thank Lim Mengyu and Reena Koh ChengYee (Nanyang Technological University) for their assistance in editing and proofreading the article, and Benedetta Corea for her support in graphical design. Explanatory icons or illustrations placed at the left side of Figure 3 have been obtained from the open-source websites Freepik and Pixabay.

## 10. Conflict of interests

The authors declare no conflict of interest. The funders had no role in the design of the study; in the collection, analyses, or interpretation of data; in the writing of the manuscript, or in the decision to publish the results.

### 11. Abbreviations

The following abbreviations are used in this manuscript:

ECR-R: Experience in Close Relationships-Revised
PBI: Prental Bonding Instrument
OXTR: Oxytocin receptor gene
SS: Sum of squares
RSS: Residual sum of squares
HMR: Hierarchical multiple regression

## 12. Ethics

The research was approved by the Ethics Committee of Nanyang Technological University (IRB-2015-08-020-01) and of University of Trento (Prot-2017-019). Informed consent was obtained from all participants, and the study was conducted following the Declaration of Helsinki. The genetic assessment was conducted on anonymized biosamples at the Nanyang Technological University (Singapore) and at the Department of Neurobiology and Behavior at Nagasaki University (Japan). Results from questionnaires were anonymized at the beginning of the collection.

## 13. Data Repository

The data of this study can be found in the NTU’s Data repository (DR-NTU Data) at the following address: https://doi.org/10.21979/N9/7NT0EG

