## Supplementary material for "Recalled parental bonding interacts with oxytocin receptor gene polymorphism in modulating anxiety and avoidance in adult relationships": https://www.mdpi.com/2076-3425/11/4/496/s1

### Supplementary Results

*Italian Sample*

*Avoidance*

Table S4 of the Supplementary Materials illustrates the series of hierarchical multiple regression conducted on ECR-R avoidance for the Italian sample (total SS = 22689). At step 1, adding OXTr to the model explained a non-significant 1% of variance ($F$(1,95) = 0.70, RSS = 22540, $p$= .41). At step 2, maternal care contributed significantly to the model and accounted for 6% of the variation in avoidance ($F$(1,94) = 6.70, RSS = 21122, $p$< .01). At step 3, the addition of maternal overprotection increased the model non-significantly by 1% ($F$(1,93) = 1.49, RSS = 20807, $p$= .23), although maternal care was still identified as positive predictor. Paternal care was entered at step 4, but no change in the $\mathbf{R}^2$ was observed ($F$(1,92) = 0.53, RSS = 20695, $p$= .47). No significant increase in the $\mathbf{R}^2$ was not even found when the variable paternal overprotection was included at step 5 ($F$(1,91) = 0.01, RSS = 20693, $p$= .93). At step 6, the hypothesized interaction between OXTr and paternal bonding subscales on avoidance was tested ($f^2$ = .22, power = .94). The final model estimates the highest proportion of explained variance in avoidance by the model ($R^2$ = .19) with a significant further 10% ($F$(4,87) = 2.68, RSS = 18422, $p$< .04). Here, the slight effect of maternal care did not reach the significance after multiple tests' correction ($\beta$ = -0.02, SE = 0.01, $t$ = -2.26, $p$< .03).

*Singaporean Sample*

*Anxiety*

Table S5 of the Supplementary Materials shows the series of hierarchical multiple regression conducted on ECR-R anxiety of the Singaporean sample (total SS = 91312). The introduction of OXTr at step 1 did not increase the explained variation of the model ($F$(1,214) = 0.38, RSS = 91158, $p$= .54). Maternal care was entered at step 2 and a not significant 1% of variance was detected ($F$(1,213) = 0.96, RSS = 90771, $p$= .33). No variation in $\mathbf{R}^2$ was found at step 3, when maternal overprotection was included ($F$(1,212) = 0.01, RSS = 90769, $p$= .95). At step 4, paternal care in input motivated a significant change in $\mathbf{R}^2$ ($F$(1,211) = 4.65, RSS = 88901, $p$= .03) and disclosed a main effect of the same PBI dimension on anxiety. At step 5, paternal overprotection was entered and accounted for a significant 2% of additional variance ($F$(1,210) = 6.35, RSS = 86349, $p$= .01). At this level, only paternal overprotection was a positive predictor of anxiety. At step 6, the hypothesized interaction between OXTr and paternal bonding dimensions on anxiety was verified ($f^2$ = .10, power = 0.95). Although most of the variance in anxiety ($R^2$ = .09) depended on the final model, no significant difference between step 6 and 5 was discovered ($F$(4,206) = 2.21, RSS = 82799, $p$= .07). Considering the application of multiple tests' correction, the interaction effect between OXTr and maternal overprotection on the anxiety levels did not reach an acceptable significance level at the final step

($\beta$ = -1.06, SE = 0.49, $t$ = -2.17, $p$= .03).

*Total Sample*

*Anxiety*

Table S6 of the Supplementary Materials reports the series of hierarchical multiple regression performed on ECR-R anxiety of the total sample (total SS = 134493). At step 1, culture was considered as starting variable of the regression model, explaining a significant 3% of the total variance ($F$(1,311) = 11.67, RSS = 130105, $p$= .0007). Culture was significant at this step, as positive predictor. When entering OXTr at step 2, no variation in $R^2$ was observed ($F$(2,309) = 0.22, RSS = 129940, $p<$ .80), but culture still obtained a significant effect. The addition of maternal care to the model at step 3 increased the variance of a significant 3% ($F$(1,308) = 9.12, RSS = 126510, $p<$ .003). Both culture and maternal care were positive predictors at this step. At step 4, although maternal overprotection improved the model by a non-significant 1% of variation in anxiety ($F$(1,307) = 2.52, RSS = 125561, $p$= .11), culture and maternal care were still the strongest predictors of the model. Paternal care contributed significantly in explaining a further 3% of variance at step 5 ($F$(1,306) = 9.63, RSS = 121937, $p$= .002). Here, culture and paternal care best predict participants' anxiety. At step 6, the parental overprotection dimension was introduced and accounted for a significant 2% of variance ($F$(1,305) = 5.01, RSS = 120053, $p$= .026). At this step, the three variable culture, paternal care and paternal overprotection resulted significant predictors. At step 7, the hypothesized interaction between culture, gene and paternal bonding dimensions on anxiety was explored ($f^2$ = .09, power = .98). The final model computed the highest proportion of explained variance in anxiety by the model ($R^2$ = .21), as highlighted by the significant increase of $R^2$ ($F$(22,283) = 1.64, RSS = 106451, $p$= .036). From the overall interactive model, no significant main effects of culture, genotype or caregiving behavior as well as two-way or three-way interactions were found for adult anxiety.

### Materials: Supplementary Tables

*Table S1: Age and Sex differences between the rs53576 genetic groups for the Italian, Singaporean and Total sample.*

| | **Age** | |
|---|---|---|
| **Sample** | **Contrast between Genotypes** | ***t*** |
| Italian | G/G - A | 1.76 (.38) |
| Singaporean | A/A - G | 0.48 (.63) |
| | A/A - A/G | -0.81 (.88) |
| Total | A/A - G/G | 0.42 (.41) |
| | G/G - A/G | -0.99 (.32) |
| | **Sex** | |
| **Sample** | **Contrast between Genotypes** | $X^2$ ***(1)*** |
| Italian | G/G - A | 0.02 (.90) |
| Singaporean | A/A - G | 0.19 (.66) |
| Total | A/A - A/G - G/G | 0.01 (.97) |

Table 1: **Age**. Statistics of Student's $t$-test on age differences between rs53576 genetic groups within the Italian (G/G vs. A-carriers), Singaporean (A/A vs. G-carriers) and the Total sample (A/A vs. A/G; A/A vs. G/G; G/G vs. A/G). **Sex**. Statistics of Pearson's $X^2$ determine the difference between the frequency of males and females belonging to each genetic group for the Italian (G/G vs. A-carriers), Singaporean (A/A vs. G-carriers) and the Total sample (A/A vs. G/G vs. A/G). For each statistical test, the $p$-value is reported between parentheses.

*Table S2: Models applied into the six-steps hierarchical multiple regressions on the Italian, Singaporean and Total sample*

| **Italian Sample** | |
|---|---|
| *Step* | **Tested Model** |
| **1** | ECR-Rvariable = *OXTR rs53576* |
| **2** | ECR-Rvariable = (*OXTR rs53576* + *M_Care*) |
| **3** | ECR-Rvariable = (*OXTR rs53576* + *M_Care* + *M_Overp*) |
| **4** | ECR-Rvariable = (*OXTR rs53576* + *M_Care* + *M_Overp* + *F_Care*) |
| **5** | ECR-Rvariable = (*OXTR rs53576* + *M_Care* + *M_Overp* + *F_Care* + *F_Overp*) |
| **6** | ECR-Rvariable = (*OXTR rs53576* * (*M_Care* + *M_Overp* + *F_Care* + *F_Overp*) |
| **Singaporean Sample** | |
| *Step* | **Tested Model** |
| **1** | ECR-Rvariable = *OXTR rs53576* |
| **2** | ECR-Rvariable = (*OXTR rs53576* + *M_Care*) |
| **3** | ECR-Rvariable = (*OXTR rs53576* + *M_Care* + *M_Overp*) |
| **4** | ECR-Rvariable = (*OXTR rs53576* + *M_Care* + *M_Overp* + *F_Care*) |
| **5** | ECR-Rvariable = (*OXTR rs53576* + *M_Care* + *M_Overp* + *F_Care* + *F_Overp*) |
| **6** | ECR-Rvariable = (*OXTR rs53576* * (*M_Care* + *M_Overp* + *F_Care* + *F_Overp*) |
| **Total Sample** | |
| *Step* | **Tested Model** |
| **1** | ECR-Rvariable = *Culture* |
| **2** | ECR-Rvariable = (*Culture* + *OXTR rs53576*) |
| **3** | ECR-Rvariable = (*Culture* + *OXTR rs53576* + *M_Care*) |
| **4** | ECR-Rvariable = (*Culture* + *OXTR rs53576* + *M_Care* + *M_Overp*) |
| **5** | ECR-Rvariable = (*Culture* + *OXTR rs53576* + *M_Care* + *M_Overp* + *F_Care*) |
| **6** | ECR-Rvariable = (*Culture* + *OXTR rs53576* + *M_Care* + *M_Overp* + *F_Care* + *F_Overp*) |
| **7** | ECR-Rvariable = (*Culture* * *OXTR rs53576* * (*M_Care* + *M_Overp* + *F_Care* + *F_Overp*) |

Table 2: As regards the Italian sample, for each ECR-R dimension (anxiety and avoidance separately) a six-steps hierarchical multiple regressions was performed with the *OXTR* gene genotype rs53576 as a between-subject factor (G/G and A-carriers) and the PBI dimensions (maternal mare, maternal overprotection, paternal care, paternal overprotection) as continuous predictors. As regards the Singaporean sample, for each ECR-R dimension (anxiety and avoidance individually) a six-steps hierarchical multiple regressions was performed with the *OXTR* gene genotype rs53576 as a between-subject factor (A/A and G-carriers) and the PBI dimensions (maternal mare, maternal overprotection, paternal care, paternal overprotection) as continuous predictors. As regards the total sample, for each ECR-R dimension (anxiety and avoidance distinctly) a seven-steps hierarchical multiple regressions was performed with the *OXTR* gene genotype rs53576 as a between-subject factor (A/A, A/G and G/G) and the PBI dimensions (maternal mare, maternal overprotection, paternal care, paternal overprotection) as continuous covariates.

*Table S3: Correlation matrix among Questionnaire Subscales across all participants*

| *subscale* | **M_Care** | **M_Overp** | **F_Care** | **F_Overp** | **Anxiety** | **Avoidance** |
|---|---|---|---|---|---|---|
| **M_Care** | | | | | | |
| **M_Overp** | -.29*** | | | | | |
| **F_Care** | .43*** | -.18* | | | | |
| **F_Overp** | -.07 | .40*** | -.20** | | | |
| **Anxiety** | -.17* | .11 | -.24*** | .15 | | |
| **Avoidance** | -.12 | .07 | -.13 | .07 | .39*** | |

Table 3: Pearson correlation coefficients and significance values among questionnaire subscales of the total sample; significance is adjusted for multiple tests (corrected *alpha* = 0.003). *$p$<.003 **$p$<.001 ***$p$<.0001

*Table S4: Hierarchical multiple regression on adult avoidance for the Italian participants.*

| *Step* | **Variable** | $\beta$ | **SE** | $t$ | $R$ | $R^2$ | $\Delta$ $R^2$ |
|---|---|---|---|---|---|---|---|
| **1** | | | | | .08 | .01 | .01 |
| | *OXTr rs53576* | -2.47 | 3.13 | -0.79 | | | |
| **2** | | | | | .26 | .07 | .06* |
| | *OXTr rs53576* | -2.33 | 3.05 | -0.77 | | | |
| | *M_Care* | -0.01* | 0.00 | -2.51 | | | |
| **3** | | | | | .29 | .08 | .01 |
| | *OXTr rs53576* | -2.26 | 3.04 | -0.74 | | | |
| | *M_Care* | -0.01* | 0.00 | -2.02 | | | |
| | *M_OverP* | 0.30 | 0.26 | 1.19 | | | |
| **4** | | | | | .30 | .09 | .01 |
| | *OXTr rs53576* | -2.07 | 3.06 | -0.68 | | | |
| | *M_Care* | -0.01 | 0.01 | -1.47 | | | |
| | *M_OverP* | 0.29 | 0.26 | 1.11 | | | |
| | *F_Care* | -0.14 | 0.19 | -0.71 | | | |
| **5** | | | | | .30 | .09 | 0 |
| | *OXTr rs53576* | -2.07 | 3.08 | -0.67 | | | |
| | *M_Care* | -0.01 | 0.01 | -1.42 | | | |
| | *M_OverP* | 0.30 | 0.28 | 1.05 | | | |
| | *F_Care* | -0.14 | 0.20 | -0.70 | | | |
| | *F_OverP* | -0.02 | 0.25 | -0.09 | | | |
| **6** | | | | | .43 | .19 | .10* |
| | *OXTr rs53576* | -18.31 | 14.51 | -1.26 | | | |
| | *M_Care* | -0.02* | 0.01 | -2.26 | | | |
| | *M_OverP* | 0.04 | 0.38 | 0.10 | | | |
| | *F_Care* | -0.21 | 0.31 | -0.69 | | | |
| | *F_OverP* | 0.37 | 0.36 | 1.05 | | | |
| | *OXTr rs53576 x M_Care* | 0.02 | 0.01 | 1.82 | | | |
| | *OXTr rs53576 x M_OverP* | 0.44 | 0.55 | 0.80 | | | |
| | *OXTr rs53576 x F_Care* | 0.29 | 0.40 | 0.72 | | | |
| | *OXTr rs53576 x F_OverP* | -0.81 | 0.50 | -1.63 | | | |

Table 4: Hierarchical multiple regression on ECR-R avoidance for the Italian sample. Note. SE =standard error of unstandardized coefficient. *$p$<.05

*Table S5: Hierarchical multiple regression on adult anxiety for the Singaporean participants.*

| *Step* | **Variable** | $\beta$ | **SE** | ***t*** | ***R*** | $\mathbf{R}^2$ | $\Delta\ \mathbf{R}^2$ |
|---|---|---|---|---|---|---|---|
| **1** | | | | | .04 | .00 | .00 |
| | *OXTr rs53576* | -1.75 | 2.91 | -0.60 | | | |
| **2** | | | | | .08 | .01 | .01 |
| | *OXTr rs53576* | -1.95 | 2.92 | -0.67 | | | |
| | *M_Care* | -0.19 | 0.20 | -0.95 | | | |
| **3** | | | | | .08 | .01 | .00 |
| | *OXTr rs53576* | -1.93 | 2.93 | -0.66 | | | |
| | *M_Care* | -0.20 | 0.21 | -0.93 | | | |
| | *M_OverP* | -0.01 | 0.21 | -0.07 | | | |
| **4** | | | | | .16 | .03 | .02* |
| | *OXTr rs53576* | -1.32 | 2.95 | -0.45 | | | |
| | *M_Care* | 0.00 | 0.23 | -0.01 | | | |
| | *M_OverP* | -0.04 | 0.21 | -0.19 | | | |
| | *F_Care* | -0.40* | 0.19 | -2.11 | | | |
| **5** | | | | | .23 | .05 | .02* |
| | *OXTr rs53576* | -1.44 | 2.89 | -0.50 | | | |
| | *M_Care* | -0.05 | 0.23 | -0.20 | | | |
| | *M_OverP* | -0.25 | 0.22 | -1.11 | | | |
| | *F_Care* | -0.33 | 0.19 | -1.75 | | | |
| | *F_OverP* | 0.57* | 0.23 | 2.49 | | | |
| **6** | | | | | .31 | .09 | .04 |
| | *OXTr rs53576* | -0.59 | 15.61 | -0.04 | | | |
| | *M_Care* | -0.66 | 0.41 | -1.60 | | | |
| | *M_OverP* | 0.54 | 0.41 | 1.32 | | | |
| | *F_Care* | 0.08 | 0.35 | 0.22 | | | |
| | *F_OverP* | 0.31 | 0.39 | 0.80 | | | |
| | *OXTr rs53576 x M_Care* | 0.81 | 0.49 | 1.64 | | | |
| | *OXTr rs53576 x M_OverP* | -1.06* | 0.49 | -2.17 | | | |
| | *OXTr rs53576 x F_Care* | -0.45 | 0.42 | -1.09 | | | |
| | *OXTr rs53576 x F_OverP* | 0.30 | 0.48 | 0.62 | | | |

Table 5: Hierarchical multiple regression on ECR-R anxiety for the Singaporean sample. Note. SE =standard error of unstandardized coefficient. *$p$<.05

*Table S6: Hierarchical multiple regression on adult anxiety for all the participants.*

| *Step* | **Variable** | $\beta$ | **SE** | $t$ | $R$ | $R^2$ | $\Delta R^2$ |
|---|---|---|---|---|---|---|---|
| **1** | | | | | .18 | .03 | .03*** |
| | *Culture* | 8.10** | 2.50 | 3.24 | | | |
| **2** | | | | | .18 | .03 | .00 |
| | *Culture* | 7.56** | 2.65 | 2.85 | | | |
| | *OXTr rs53576 A/G-A/A* | -0.64 | 2.80 | -0.23 | | | |
| | *OXTr rs53576 G/G-A/A* | -1.96 | 3.19 | -0.61 | | | |
| **3** | | | | | .24 | .06 | .03** |
| | *Culture* | 7.01** | 2.63 | 2.67 | | | |
| | *OXTr rs53576 A/G-A/A* | -1.02 | 2.77 | -0.37 | | | |
| | *OXTr rs53576 G/G-A/A* | -2.48 | 3.16 | -0.78 | | | |
| | *M_Care* | -0.44** | 0.15 | -2.89 | | | |
| **4** | | | | | .26 | .07 | .01 |
| | *Culture* | 7.27** | 2.63 | 2.77 | | | |
| | *OXTr rs53576 A/G-A/A* | -1.22 | 2.77 | -0.44 | | | |
| | *OXTr rs53576 G/G-A/A* | -3.04 | 3.17 | -0.96 | | | |
| | *M_Care* | -0.37* | 0.16 | -2.32 | | | |
| | *M_OverP* | 0.27 | 0.18 | 1.52 | | | |
| **5** | | | | | .31 | .09 | .03** |
| | *Culture* | 6.90** | 2.60 | 2.66 | | | |
| | *OXTr rs53576 A/G-A/A* | -0.47 | 2.74 | -0.17 | | | |
| | *OXTr rs53576 G/G-A/A* | -1.90 | 3.16 | -0.60 | | | |
| | *M_Care* | -0.16 | 0.17 | -0.93 | | | |
| | *M_OverP* | 0.23 | 0.18 | 1.29 | | | |
| | *F_Care* | -0.45** | 0.15 | -3.02 | | | |
| **6** | | | | | .33 | .11 | .02* |
| | *Culture* | 7.82** | 2.61 | 2.99 | | | |
| | *OXTr rs53576 A/G-A/A* | -0.18 | 2.73 | -0.07 | | | |
| | *OXTr rs53576 G/G-A/A* | -1.77 | 3.14 | -0.56 | | | |
| | *M_Care* | -0.21 | 0.17 | -1.19 | | | |
| | *M_OverP* | 0.07 | 0.19 | 0.37 | | | |
| | *F_Care* | -0.39* | 0.15 | -2.55 | | | |
| | *F_OverP* | 0.41* | 0.19 | 2.19 | | | |
| **7** | | | | | .46 | .21 | .10* |
| | *Culture* | 36.09 | 33.32 | 1.08 | | | |
| | *OXTr rs53576 A/G-A/A* | 20.05 | 35.92 | 0.56 | | | |
| | *OXTr rs53576 G/G-A/A* | 37.17 | 34.54 | 1.08 | | | |
| | *M_Care* | -0.01 | 1.02 | -0.02 | | | |
| | *M_OverP* | 4.00 | 2.05 | 1.95 | | | |
| | *F_Care* | -1.45 | 0.89 | -1.62 | | | |
| | *F_OverP* | -0.53 | 1.05 | -0.50 | | | |
| | *Culture x OXTr rs53576 A/G-A/A* | -17.77 | 39.35 | -0.45 | | | |
| | *Culture x OXTr rs53576 G/G-A/A* | -46.96 | 40.14 | -1.17 | | | |
| | *Culture x M_Care* | -0.64 | 1.09 | -0.59 | | | |
| | *Culture x M_OverP* | -3.46 | 2.09 | -1.66 | | | |
| | *Culture x F_Care* | 1.53 | 0.95 | 1.60 | | | |
| | *Culture x F_OverP* | 0.84 | 1.12 | 0.76 | | | |
| | *OXTr rs53576 A/G-A/A x M_Care* | -0.05 | 1.16 | -0.04 | | | |
| | *OXTr rs53576 G/G-A/A x M_Care* | -0.42 | 1.07 | -0.39 | | | |
| | *OXTr rs53576 A/G-A/A x M_OverP* | -2.87 | 2.15 | -1.34 | | | |
| | *OXTr rs53576 G/G-A/A x M_OverP* | -3.12 | 2.11 | -1.48 | | | |
| | *OXTr rs53576 A/G-A/A x F_Care* | 0.83 | 1.03 | 0.81 | | | |
| | *OXTr rs53576 G/G-A/A x F_Care* | 0.94 | 0.95 | 0.98 | | | |
| | *OXTr rs53576 A/G-A/A x F_OverP* | 0.36 | 1.27 | 0.28 | | | |
| | *OXTr rs53576 G/G-A/A x F_OverP* | 0.28 | 1.15 | 0.24 | | | |
| | *Culture x OXTr rs53576 A/G-A/A x M_Care* | 1.07 | 1.27 | 0.84 | | | |
| | *Culture x OXTr rs53576 G/G-A/A x M_Care* | 0.93 | 1.23 | 0.75 | | | |
| | *Culture x OXTr rs53576 A/G-A/A x M_OverP* | 1.57 | 2.20 | 0.71 | | | |
| | *Culture x OXTr rs53576 G/G-A/A x M_OverP* | 2.77 | 2.21 | 1.25 | | | |
| | *Culture x OXTr rs53576 A/G-A/A x F_Care* | -1.54 | 1.11 | -1.38 | | | |
| | *Culture x OXTr rs53576 G/G-A/A x F_Care* | -1.03 | 1.08 | -0.95 | | | |
| | *Culture x OXTr rs53576 A/G-A/A x F_OverP* | 0.07 | 1.36 | 0.05 | | | |
| | *Culture x OXTr rs53576 G/G-A/A x F_OverP* | -0.49 | 1.32 | -0.37 | | | |

Table 6: Hierarchical multiple regression on ECR-R anxiety for the total sample. Note. SE =standard error of unstandardized coefficient. *$p$<.05 **$p$<.01 ***$p$<.001

*Table S7: Means and Standard Errors of ECR-R anxiety by the independent variable for the Italian participants*

| **PBI Dimension** | **Low** | **High** |
|---|---|---|
| Maternal Overprotection | 52.41 (2.18) | 62.07 (3.58) |

Table 7: Means and Standard Error values in low and high maternal overprotection on ECR-R anxiety for the main effect observed at the final step of the hierarchical multiple regressions on the Italian sample. Standard Error Means (SEM) are reported between parentheses.

*Table S8: Means and Standard Errors of ECR-R avoidance by the independent variables for Singaporean participants*

| **PBI Dimension** | **Low** | **High** | | |
|---|---|---|---|---|
| A. Maternal Overprotection | 41.21 (1.57) | 39.40 (1.73) | | |
| **PBI Dimension** | **Low/AA** | **Low/G** | **High/AA** | **High/G** |
| B: Maternal Overprotection | 43.47 (3.00) | 40.10 (1.83) | 36.55 (2.71) | 41.38 (2.22) |

Table 8: Means and Standard Error values of ECR-R avoidance for each significant main and interaction effect confirmed by the final step of the hierarchical multiple regressions on the Singaporean sample. **A**. Mean values in low and high maternal overprotection on avoidance. **B**. Mean values in A/A homozygotes and G-carriers divided in low and high maternal overprotection on avoidance.

*Table S9: Means and Standard Errors of ECR-R avoidance by the independent variables for all the participants assessed*

| **PBI Dimension** | **Low** | **High** | | | | |
|---|---|---|---|---|---|---|
| A. Maternal Overprotection | 37.56 (1.30) | 39.96 (1.38) | | | | |
| **PBI Dimension** | **Low/Italian** | **Low/Singaporean** | **High/Italian** | **High/Singaporean** | | |
| B. Maternal Overprotection | 31.63 (2.02) | 39.87 (1.59) | 38.28 (2.28) | 40.84 (1.73) | | |
| **PBI Dimension** | **Low/AA** | **Low/AG** | **Low/GG** | **High/AA** | **High/AG** | **High/GG** |
| C. Maternal Overprotection | 37.40 (2.57) | 40.36 (1.82) | 32.95 (2.39) | 2.68 (0.05) | 37.72 (2.35) | 40.62 (2.12) |
| **PBI Dimension** | **Italian/AA** | **Italian/AG** | **Italian/GG** | **Singaporean/AA** | **Singaporean/AG** | **Singaporean/GG** |
| D. Low Maternal Overprotection | 29.50 (6.40) | 32.71 (2.81) | 31.71 (2.89) | 38.75 (2.78) | 42.31 (2.10) | 34.81 (4.18) |
| E. High Maternal Overprotection | 51.33 (4.60) | 37.10 (3.78) | 36.04 (3.13) | 40.97 (2.99) | 38.05 (3.02) | 44.85 (2.67) |

Table 9: Means and Standard Error values of ECR-R avoidance for each significant main and interaction effect obtained from the final step of the hierarchical multiple regressions on the total sample. **A**. Mean values in low and high maternal overprotection on avoidance. **B**. Mean values in the Italian and Singaporean participants divided in low and high maternal overprotection on avoidance. **C** Mean values in A/A homozygotes, A/G heterozygotes and G/G homozygotes divided in low and high maternal overprotection on avoidance. **D** Mean values in A/A homozygotes, A/G heterozygotes and G/G homozygotes divided in Italian and Singaporean participants with low maternal oveprotection on avoidance. **E** Mean values in A/A homozygotes, A/G heterozygotes and G/G homozygotes divided in Italian and Singaporean participants with high maternal oveprotection on avoidance. Standard Error Means (SEM) are reported between parentheses.

*Figure S1: Comparison between levels of sex and age on anxiety and avoidance for the Italian participants*

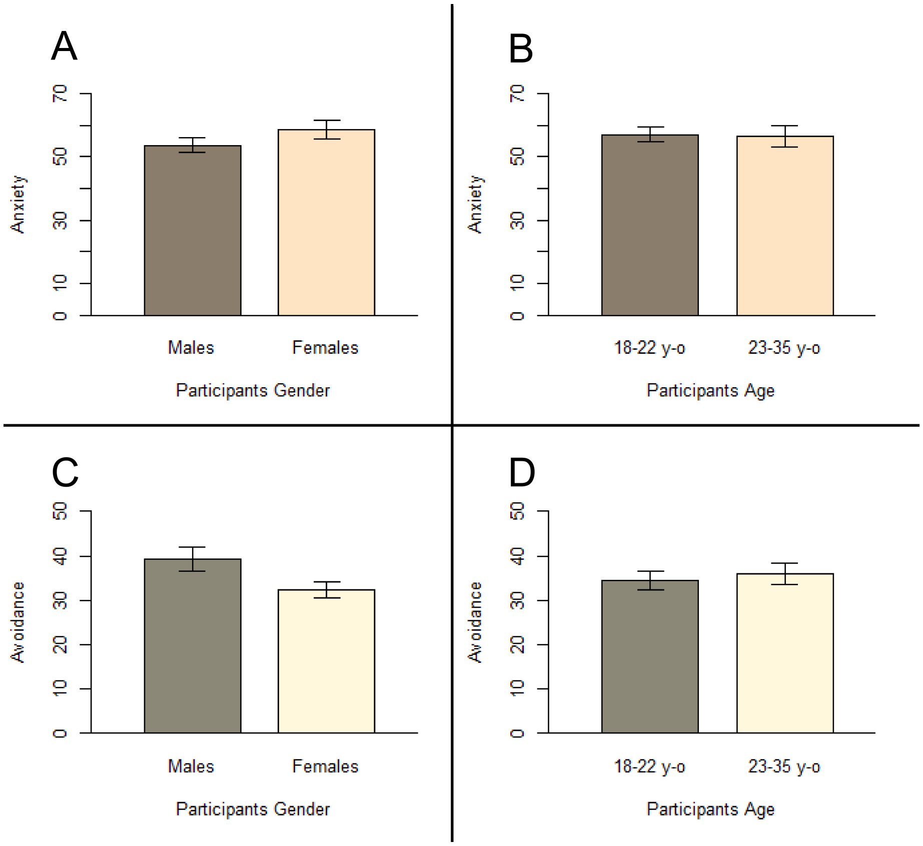


Figure 1: (**A**). Contrast between Italian male and Italian female participants on ECR-R anxiety. (**B**). Contrast between Italian participants aged between 18 and 22 years-old and Italian participants aged between 23 and 35 years-old on ECR-R anxiety. (**C**). Contrast between Italian male and Italian female participants on ECR-R avoidance. (**D**). Contrast between Italian participants aged between 18 and 22 years-old and Italian participants aged between 23 and 35 years-old on ECR-R avoidance. Median split procedure was applied to convert the continuous variable age to a two-levels factor, as visible from **B** and **C**. No significant differences were found.

*Figure S2: Comparison between levels of sex and age on anxiety and avoidance for the Singaporean participants*

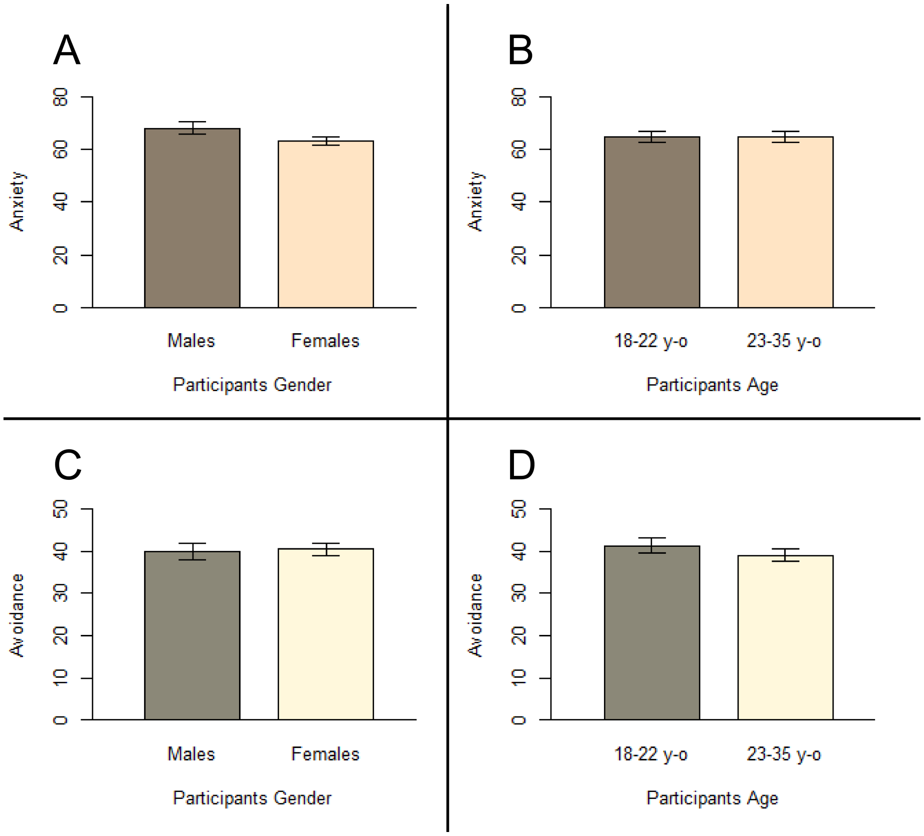


Figure 2: (**A**). Contrast between Singaporean male and Singaporean female participants on ECR-R anxiety. (**B**). Contrast between Singaporean participants aged between 18 and 22 years-old and Singaporean participants aged between 23 and 35 years-old on ECR-R anxiety. (**C**). Contrast between Singaporean male and Singaporean female participants on ECR-R avoidance. (**D**). Contrast between Singaporean participants aged between 18 and 22 years-old and Singaporean participants aged between 23 and 35 years-old on ECR-R avoidance. Median split procedure was applied to convert the continuous variable age to a two-levels factor, as visible from **B** and **C**. No significant differences were found.

*Figure S3: Comparison between levels of sex, age and culture on anxiety and avoidance for all the participants assessed*

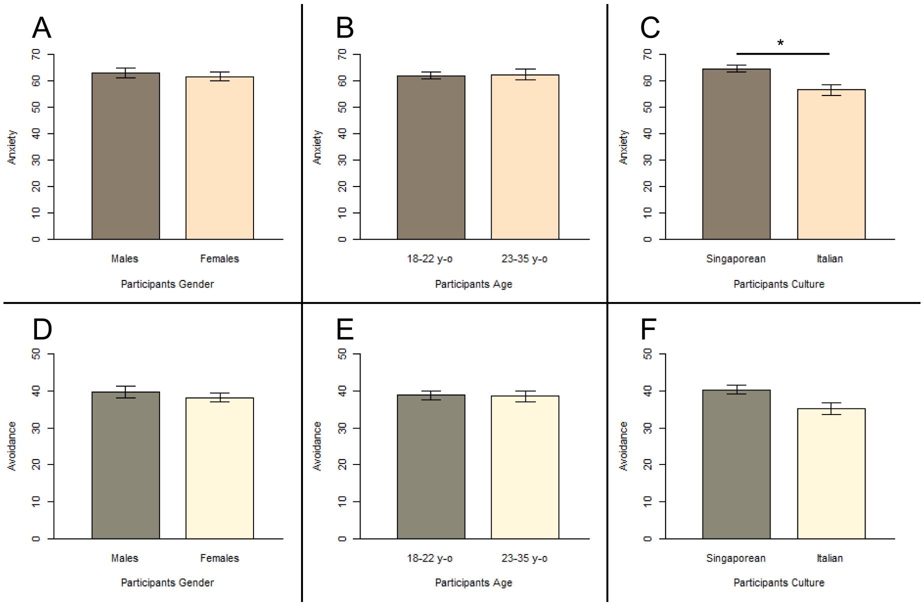


Figure 3: (**A**). Contrast between male and female participants on ECR-R anxiety. (**B**). Contrast between participants aged between 18 and 22 years-old and participants aged between 23 and 35 years-old on ECR-R anxiety. **C**). Effect of culture on ECR-R anxiety. Contrast between Singaporean and Italian participants on ECR-R anxiety. (**D**). Contrast between male and female participants on ECR-R avoidance. (**E**). Contrast between participants aged between 18 and 22 years-old and participants aged between 23 and 35 years-old on ECR-R avoidance. **F**). Contrast between Singaporean and Italian participants on ECR-R avoidance. Median split procedure was applied to convert the continuous variable age to a two-levels factor, as visible from **B** and **E**. **D**). * p ¡ 0.008
